# Microbial community profiles of genital and non-genital skin lesions from Ghana

**DOI:** 10.64898/2026.10.06.757009

**Authors:** Yaw Agyekum Boaitey, Alex Owusu-Ofori, Philipp P. Bosshard, Farhang Aghakhanian, Christopher M. Hennelly, Saki Raheem, Natasha Arora, Jonathan B. Parr, Pascale Gerbault

## Abstract

Skin ulcers, such as those observed in cases of the non-venereal disease yaws and the venereal disease syphilis, are traditionally diagnosed based on their anatomical location and visible appearance. Yet various evidence suggests that the pathogens that cause these ulcers may not be restricted to genital or non-genital body sites. We explored this by profiling the bacterial communities of genital and non-genital skin ulcers collected in Ghana using 16S rRNA gene amplicon sequencing.

Swab samples were collected from 156 clinically diagnosed skin lesions (110 non-genital and 46 genital). The V1–V2 region of the 16S rRNA gene was sequenced to characterize bacterial communities. Microbial diversity, community composition, differential abundance, and associations between lesion diversity and common skin bacteria were analysed using bioinformatics and statistical methods.

Following quality filtering, 135 samples were retained, revealing 23 bacterial phyla, 143 families, and 406 genera. Alpha diversity indicated similar bacterial richness and diversity in genital and non-genital skin lesions. Beta diversity analyses indicated differences between genital and non-genital lesions, accompanied by significant differences in within group dispersion. A log2 fold change analysis evidenced genital lesions were enriched in *Gardnerella*, whereas non-genital lesions showed more abundance of *Staphylococcus*, *Streptococcus*, *Corynebacterium*, *Bradyrhizobium*, and *Pelomonas*. Clinically important genera, including *Treponema*, *Haemophilus*, and *Mycobacterium*, were also detected, even in samples with low sequencing depths. Statistical modelling indicated that greater microbial diversity was associated with intermediate abundances of common skin taxa, particularly *Corynebacteriaceae*, *Micrococcaceae*, *Cutibacterium*, and *Staphylococcaceae*, thereby attesting the presence of polymicrobial communities in skin ulcers.

Overall, genital and non-genital skin lesions showed similar microbial diversity but differed in composition. Our findings support the view of skin lesions as ecological niches containing diverse bacterial communities rather than being consistently dominated by a single bacterial taxon. They provide a basis for investigating possible polymicrobial interactions in skin ulcers.

## Introduction

Skin lesions have numerous causes and are often classified according to anatomical location and clinical presentation (1, 2). Skin diseases affect nearly one-third of the global population (3) and include chronic inflammatory disorders, malignancies, genetic conditions, and a wide range of bacterial, viral, fungal, and parasitic infections (4). In tropical regions of Africa and the South Pacific, infectious skin lesions are estimated to affect about 100,000 children each year (5, 6). In response to the heavy burden they inflict in those regions, the World Health Organization (7) has introduced an integrated framework for the control and management of skin-related neglected tropical diseases (skin NTDs).

Yaws, a skin ulcer disease primarily affecting children in tropical regions, is one such condition (2, 8, 9). The disease is caused by the bacterial pathogen *Treponema pallidum* subsp. *pertenue* (2, 10). Diagnosis of yaws in endemic regions often relies on clinical assessment, including anatomical location of skin ulcers due to limited laboratory capacity (2, 9, 11). However, this approach is unreliable for several reasons. On the one hand, several pathogens, including *Mycobacterium ulcerans* (Buruli ulcer) may cause skin ulcers of similar clinical appearance (2, 12–17). On the other hand, molecular techniques have demonstrated that lesions clinically diagnosed as yaws may host other pathogens, including *M. ulcerans* and *Haemophilus ducreyi* (typically associated with genital chancroid), challenging the assumption that ulcer location reliably predicts disease aetiology (3, 15, 18–20). In addition to these issues, analyses of the microbial communities in skin ulcers by next-generation sequencing (NGS) including 16S rRNA amplicon sequencing have provided broader profiling of bacterial communities (6, 21–24). They have challenged the assumption that a single pathogen causes ulcer formation.

Similar diagnostic challenges occur for lesions found in the genital area, often considered to be sexually transmitted infections, and generally classified as genital ulcer disease (GUD). In fact, non-sexually acquired bacterial, skin, and fungal infections, or illnesses from non-infectious causes (e.g. fixed drug eruption, Behcet disease, sexual trauma), can present with similar ulcers in the genital area (1). Hence, ulcers found on the genitalia can be caused by sexually and non-sexually transmitted pathogens, making identification of the causal pathogen by clinical presentation alone challenging (1, 25). Importantly, pathogens associated with GUD, including *T. pallidum* subsp. *pallidum* and *H. ducreyi*, have also been detected in extra-genital sites, such as in anal and oropharyngeal ulcers (1, 26).

Thus, pathogens traditionally associated with GUD have also been identified in non-genital lesions, while more than one organism linked to tropical skin ulcers have been detected in clinically similar lesions across various body sites (1, 2, 26). Together, these observations suggest that ulcer-associated bacterial communities may not be strictly defined by anatomical location alone. Despite these overlaps, GUD, non-genital tropical ulcers, and skin NTDs are typically investigated separately, with limited attention to potential similarities in their broader microbial composition (1–3). Understanding whether ulcers from different anatomical sites harbor distinct or overlapping microbial communities may improve interpretation of ulcer aetiology, refine diagnostic approaches – currently relying heavily on lesion location and clinical appearance –, and provide insight into the possible contribution of polymicrobial interactions to ulcer pathology. In this study, we addressed this by comparing the bacterial community profiles of genital and non-genital skin ulcers in Ghana, a country where the skin NTD yaws is endemic. Using 16S rRNA next-generation sequencing, we characterized the microbial community structure across a range of anatomical sites and determined whether ulcer location is associated with distinct or shared microbial signatures.

## Methods

### Ethical Clearance

The study protocol was reviewed and approved by the ethical committees of the Komfo Anokye Teaching Hospital in Ghana (approval number: KATH-IRB/AP/048/20) and the School of Life Sciences, University of Westminster in London, England (approval number: ETH2021-0285). Participants were recruited from the Ashanti region of Ghana between February and December 2021. The study design, protocol and earlier analyses of the samples collected have been detailed previously (26). They are summarized below.

### Study communities

Skin lesion samples were collected as part of a study investigating the presence of *T. pallidum* subspecies in yaws-like and syphilis-like skin ulcers in Ghana (26). Yaws primarily affects children ≤15 years of age, with peak incidence occurring among those aged 6-10 years, particularly in deprived rural areas. Children presenting with yaws-like (non-genital) lesions were enrolled from six communities across three different municipalities in the Ashanti region (see Boaitey et al. (26) for further details): the Afigya Kwabre South Municipality (Adweratia and Mpobi); the Juaben Municipality (Kotei and Atia); and the Atwima Kwanwoma Municipality (Brofoyedu and Bebu).

Syphilis, in contrast, is more commonly reported among adolescents and adults, who in Ghana typically seek care at hospitals or pharmacies. Participants presenting with syphilis-like (genital) lesions were recruited from two hospitals in Kumasi, the capital of Ashanti region: Komfo Anokye Teaching Hospital (KATH) and South Suntreso Hospital (SSH).

### Community entering and data collection

The district health directorate, assembly members and chiefs were initially consulted and given copies of the protocol before recruitment. They were subsequently involved in community sensitization of yaws. Study objectives, procedures, risks and benefits were explained in Twi where needed to household heads, participants and parents or guardians, with opportunities for questions throughout.

Adults (≥18 years) with at least one lesion consistent with primary or secondary syphilis were enrolled after consenting. Written consent was obtained from literate participants while non-literate participants gave a thumbprint in the presence of an impartial witness.

Children aged 5–17 years with at least one lesion consistent with primary or secondary yaws were enrolled after giving verbal assent, supported by a pictorial information sheet, with written consent from a parent or guardian.

Community of residence, sex and age were recorded but sexual behaviour was outside the original study’s scope and was not collected (26). No personally identifiable information was recorded, and all participants received standard-of-care treatment per local guidelines regardless of enrolment.

### Clinical diagnosis and confirmation of treponemal infection

A total of 110 yaws-like and 46 syphilis-like lesions were clinically diagnosed by the study clinician and a trained community nurse based on the presence of lesions consistent with primary or secondary treponemal infection. Serological assessment for treponemal infection was performed using the Chembio Dual Path Platform (DPP) Screen and Confirm test (Chembio Diagnostic Systems Inc. Medford, New York) according to the manufacturer’s guidelines. This point-of-care test simultaneously detects treponemal and non-treponemal antibodies on a single platform and provides results within 15 minutes.

### Multiplex PCR

The molecular identification of *T. pallidum*, herpes simplex virus (HSV) −1 and −2 and *H. ducreyi* was achieved with a multiplex PCR protocol previously developed for routine clinical practices. This was performed at the University Hospital Zurich, Switzerland, with the Roche LightCycler 96 (Roche Diagnosis, Switzerland). This procedure specifically targets the 16S ribosomal RNA gene for both *T. pallidum* and *H. ducreyi*, and glycoprotein B (gB) region of HSV-1/-2 as described by Glatz and colleagues (27) with few alterations, outlined in Boaitey et al. (26).

### 16S ribosomal RNA gene amplification and sequencing

One moist skin lesion per participant was sampled using Dacron swabs according to a standard protocol. Skin lesion samples were then stored at −20 °C at the Kumasi Centre for Collaborative Research in Kumasi (KCCR), where DNA was extracted using QIAmp DNA mini kits (Qiagen, Germany) according to the manufacturer’s instructions. Library preparation targeted the V1-V2 variable regions of the 16S rRNA gene (ca. 300 bp), using the quick 16S Next Generation library preparation kits (Zymo research, USA) according to manufacturer’s instructions (described in the Supporting Information file **SI1** and **S1 – S10 Fig.**). A negative control was included to ensure that amplicons generated from the library preparation steps were not due to reagent contamination. Paired-end sequencing (2 x 250 bp) was carried out on an Illumina MiSeq platform (Illumina, Inc., Hayward CA, USA). Primer trimming, quality filtering, denoising and paired-end read merging and chimera removal were performed. The sequencing data is available in the NCBI Sequence Read Archive (SRA) under BioProject accession number PRJNA1530576. Paired-end reads were processed using the **DADA2** package (version 1.26.0) in R (version 4.2.2). The resulting amplicon sequence variants (ASVs) underwent taxonomic classification using Silva reference database (version 138.1) in **DADA2**. Further details are provided in the Supporting Information files **SI1**, **SI2**, **SI3**.

### Statistical analyses

Exploratory analyses and analyses of within-sample (alpha) and between-sample (beta) diversity, and taxonomic composition were conducted in R (version 4.2.2), using various packages including **vegan**, **phyloseq**, **DESeq2** and **stats** (**SI1**, **S11-S13 Fig.**). Alpha diversity was measured using observed richness, Shannon, and Simpson indices. Differences in alpha diversity between genital and non-genital lesions were assessed using Wilcoxon tests, while differences among anatomical sites were assessed using Kruskal–Wallis tests. Beta diversity was assessed using Bray–Curtis and Jaccard distances and visualized using principal coordinates analysis (PCoA). Differences in community composition were evaluated with PERMANOVA, and differences in multivariate dispersion were also evaluated. Differential abundance between genital and non-genital lesions was analyzed with **DESeq2**. Correction for multiple testing was performed using the Benjamini–Hochberg (28) method. Further details are provided in SI1. Figures were generated with **ggplot2** (version 3.4.0).

Association between microbial diversity of genital and non-genital skin lesions with relative abundance of bacterial taxa commonly found in the human skin microbiome was investigated. The taxa of interest included ASVs assigned to the phyla Actinobacteria, Bacteroidetes, Firmicutes and Proteobacteria. These taxa are referred to as “ubiquitous taxa” of the healthy skin. Within these groups, taxa frequently associated with skin microbiota, such as the families Corynebacteriaceae, Micrococcaceae, and Staphylococcaceae, as well as the genera *Cutibacterium*, were specifically targeted. The additional phyla Spirochaetes and genera *Mycobacterium* and *Haemophilus* were included into the number of taxa of interest as these include pathogenic bacteria that were identified in some of the skin lesions sampled. For each sample, the number of reads assigned to each taxon was recorded and converted into relative abundance. Associations between the relative abundance of these taxa and alpha diversity measures were assessed using linear models performed in R. Details about model comparisons, linear model diagnostics, and coefficient estimations are provided in the Supporting Information file **SI1**, **S14-S24 Fig.** and **S Tables 1-5**.

## Results

### Common pathogens in non-genital and genital skin ulcers in Ghana

Across the 156 samples analysed, a total of 3,490,536 reads were observed. Sequencing depths varied widely, ranging from 2 to 71,329 reads per sample, with a median of 18,712 and a mean of 22,375 reads per sample (**SI1**, **S11 Fig.**).

The presence of pathogens known to be associated with genital and non-genital skin lesions in Ghana (i.e. *T. pallidum* spp., *H. ducreyi* and *M. ulcerans*) was specifically investigated by examining the presence and number of *Treponema*, *Haemophilus* and *Mycobacterium* ASVs (14, 15, 20, 29, 30). Across the 156 samples analyzed, reads pertaining to these three pathogens were found in 22 samples (summarized in **Table 1**): *Treponema* reads were detected in three samples, including one from a genital lesion; *Haemophilus* reads were found in 15 samples, seven of which were from genital lesions; and *Mycobacterium* reads were detected in four samples, all from non-genital lesions.

**Table 1.** Number of 16S reads attributed to three of the common pathogens known to cause skin infectious diseases.

| Sample ID | Lesion location | DPP | qPCR | 16S Genus | nb reads | tot reads |
| --- | --- | --- | --- | --- | --- | --- |
| S018 | genital | Negative | HSV2 | *Treponema* | 9 | 29,419 |
| S022* | genital | Negative | HSV2 | *Haemophilus* | 101 | 555 |
| S023 | genital | Negative | HSV2 | *Haemophilus* | 2 | 9,768 |
| S029 | genital | Negative | HSV2 | *Haemophilus* | 8 | 8,957 |
| S033 | genital | Negative | HSV2 | *Haemophilus* | 5 | 12,068 |
| S025 | genital | Negative | - | *Haemophilus* | 5 | 10,324 |
| S027 | genital | Negative | - | *Haemophilus* | 278 | 22,691 |
| S042 | genital | Negative | - | *Haemophilus* | 244 | 11,708 |
| S020* | arm | Positive | - | *Haemophilus* | 12 | 828 |
| Y008 | arm | Negative | HSV1 | *Haemophilus* | 292 | 25,329 |
| Y064 | leg | Negative | *H. ducreyi* | *Mycobacterium* | 98 | 10,661 |
| Y029 | leg | Positive | - | *Treponema* | 3 | 35,420 |
| Y048 | arm | Negative | - | *Treponema* | 2,708 | 11,030 |
| Y110 | leg | Negative | *H. ducreyi* | *Haemophilus* | 413 | 38,609 |
| Y022* | arm | Negative | - | *Haemophilus* | 68 | 481 |
| Y041 | leg | Negative | - | *Haemophilus* | 6,127 | 9,080 |
| Y045 | leg | Negative | - | *Haemophilus* | 2 | 23,009 |
| Y071 | leg | Negative | - | *Haemophilus* | 44 | 17,941 |
| Y082 | leg | Negative | - | *Haemophilus* | 560 | 24,695 |
| Y085 | leg | Negative | - | *Mycobacterium* | 69 | 14,274 |
| Y007 | oral | Negative | - | *Mycobacterium* | 77 | 31,187 |
| Y056 | oral | Negative | - | *Mycobacterium* | 12 | 33,692 |

For these 22 samples, the results from serological testing using the DPP assay and pathogen-specific quantitative PCR (qPCR) targeting *H. ducreyi*, *T. pallidum* spp., herpes simplex virus −1 (HSV1) and −2 (HSV2) were also examined (**Table 1**). Among the eight genital lesions, HSV-2 was detected by qPCR in five samples, co-detected with either *Treponema* or *Haemophilus* 16S ASVs. The remaining genital lesions were characterized by the presence of *Haemophilus* ASVs only, with no evidence of the other targeted pathogens by qPCR or DPP testing (**Table 1**). In the 14 non-genital lesions, several samples showed evidence of more than one ulcer-associated pathogen. Co-detection patterns included *Treponema* with *Haemophilus* ASVs, HSV-1 with *Haemophilus* ASVs, and *H. ducreyi* qPCR positivity together with *Mycobacterium* ASVs.

In two cases (Y029 and Y110), pathogen detection was supported by multiple approaches, particularly concordance between qPCR or DPP testing and 16S sequencing results (**Table 1**). For the yaws-suspected leg lesion Y029, positive DPP serological test and *Treponema* 16S ASVs support a treponematosis diagnosis. The extremely low *Treponema* read abundance (<0.0001) explains the negative qPCR and likely stems from sample degradation (31–33) rather than contamination, which was absent in controls. Though species identification was impossible, serological cross-reactivity with *T. carateum* remains a possibility, while oral commensals are topographically unlikely. For the yaws-suspected sample Y110, a negative DPP serological test was consistent with the absence of *Treponema* DNA. Instead, qPCR and 16S ASVs independently identified *H. ducreyi* and *Haemophilus* sp., respectively, corroborating the established role of *H. ducreyi* as a causative agent of non-genital, yaws-like lesions (14, 15, 20, 34, 35).

Overall, these findings demonstrate that different diagnostic approaches identify overlapping but not always identical pathogen profiles. This highlights the value of combining serological and molecular methods to better characterize skin ulcer aetiology in tropical settings.

Lesion location: body site the lesion was sampled from; DPP: dual platform treponemal serological test, “Positive” for a serologically positive test - detection of treponemal antibody, “Negative” otherwise; qPCR: quantitative PCR amplification for the following targeted pathogens: *H. ducreyi*, *T. pallidum* spp., herpes simplex virus −1 (HSV1) and −2 (HSV2), or “-“ when none of these four pathogens was amplified from the corresponding lesion sample; 16S Genus: genus of interest from the 16S data from the corresponding sample, with associated number of reads (nb reads); tot reads: total number of reads in the sample, samples marked with an asterisk (*) in the Sample ID column are those with fewer than 1,000 reads in total and were excluded from downstream analyses.

### Phyla in non-genital (yaws-like) and genital (syphilis-like) lesions

A total of 21 samples (eight genital and 13 non-genital lesions) with fewer than 1,000 reads were excluded from downstream diversity analyses, including relative abundance, differential abundance, beta and alpha diversity. Across the 135 samples retained, after removing unidentified phylum sequences, a total of 23 phyla, 143 families and 406 different genera were identified (**Fig. 1**). The taxonomic barplots show that, despite apparent differences in relative abundances, similar phyla are present in both genital and non-genital lesions (**Fig. 1** and **SI1**, **S12** and **S13 Fig.**).

**Fig. 1.**
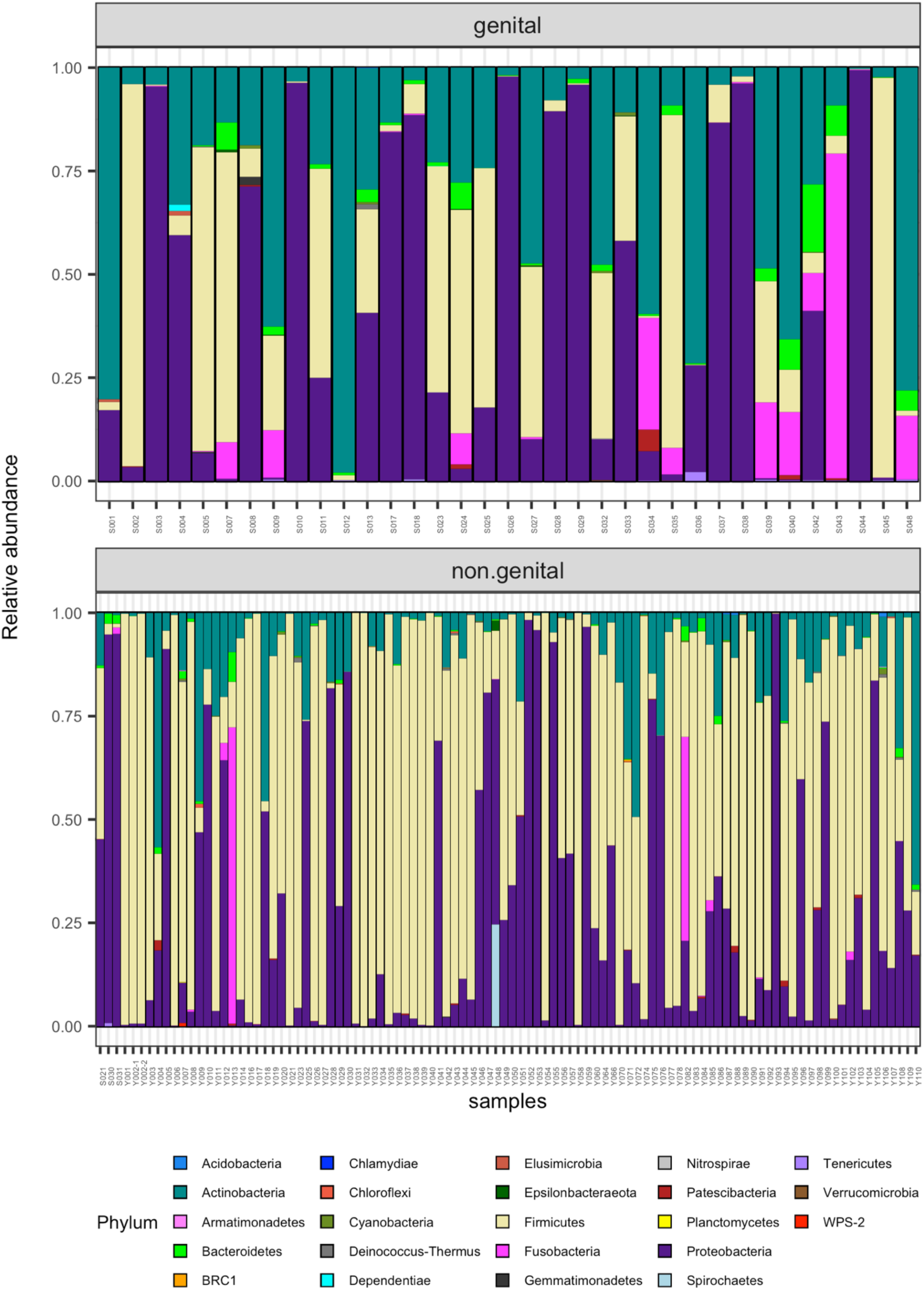
Relative abundance of bacterial phyla in genital and non-genital of 135 samples. Stacked bar charts show the composition of bacterial communities at the phylum level across individual skin lesion samples classified as genital (top panel) and non-genital (bottom panel). Each of the vertical bar represents one sample, and the color-coded segments indicate the relative abundance of different bacterial phyla, as specified in the legend. The y-axis denotes relative abundance (0–1). The same phyla (23) were present in both types of lesions, but their relative abundance differs.

### Comparison of the diversity of genital and non-genital skin ulcer microbiota

Alpha (intra sample variation) and beta diversity (inter sample variation) analyses were performed to compare community richness and dissimilarity, respectively, between genital and non-genital lesions. Alpha diversity (Observed richness, Shannon and Simpson indices) was not significantly different between genital and non-genital lesions (adjusted p-value > 0.05; **Fig. 2**). When comparing individual anatomical locations, **Fig. 3** illustrates distributions of alpha diversity values vary, but do not significantly differ (adjusted p value > 0.05).

**Fig. 2.**
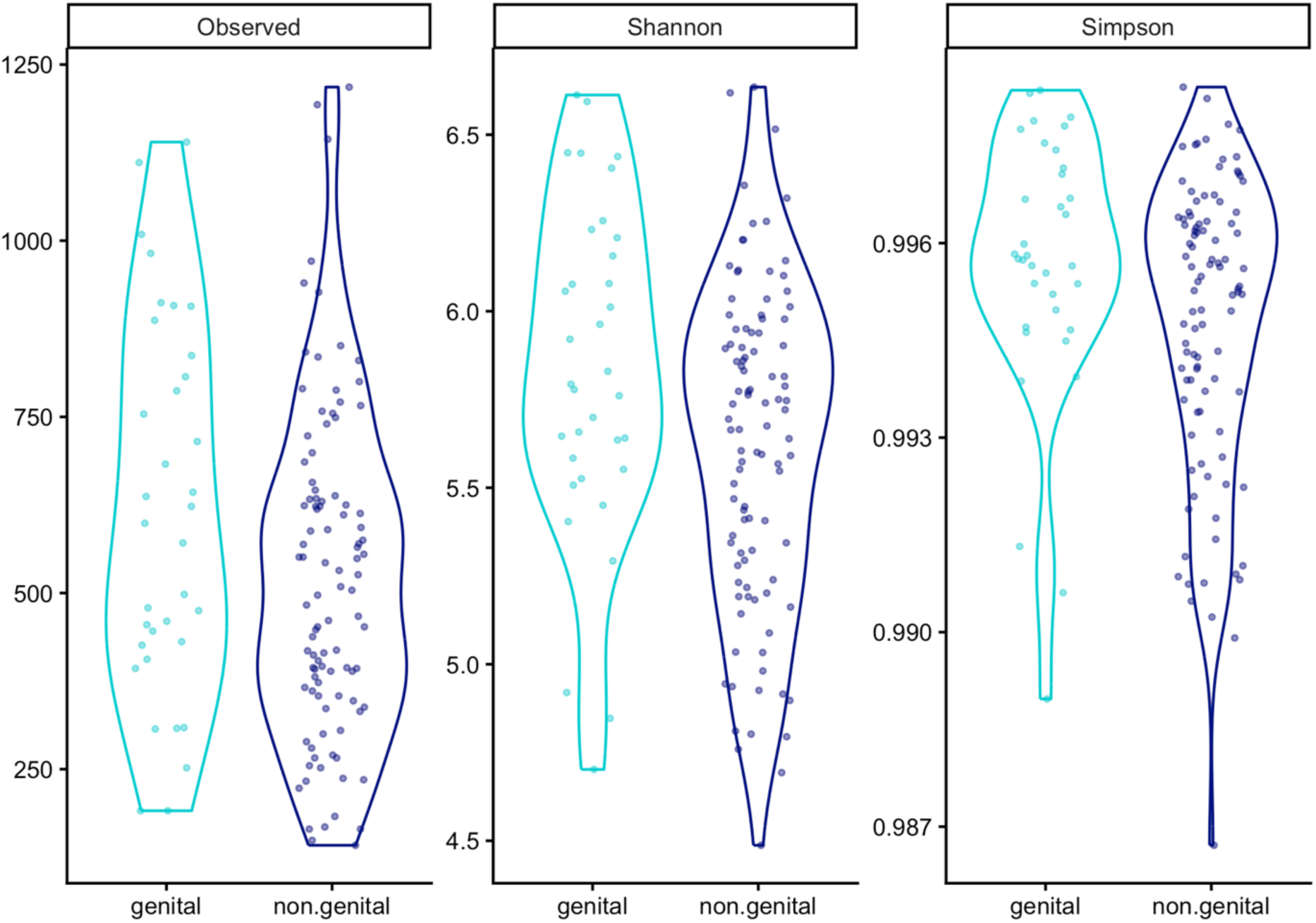
Comparison of alpha diversity between genital and non-genital sample groups. Observed richness, Shannon and Simpson diversity indices were calculated for genital and non-genital skin lesion samples. Alpha diversity distributions were statistically compared between genital and non-genital skin lesion samples with a Wilcoxon test (*wilcox.test* function in R) and a Benjamini-Hochberg correction for multiple testing was applied (*p.adjust* function with method = “BH” in R). Alpha diversity does not significantly differ between genital and non-genital skin lesion samples (adjusted p-values > 0.05).

**Fig. 3.**
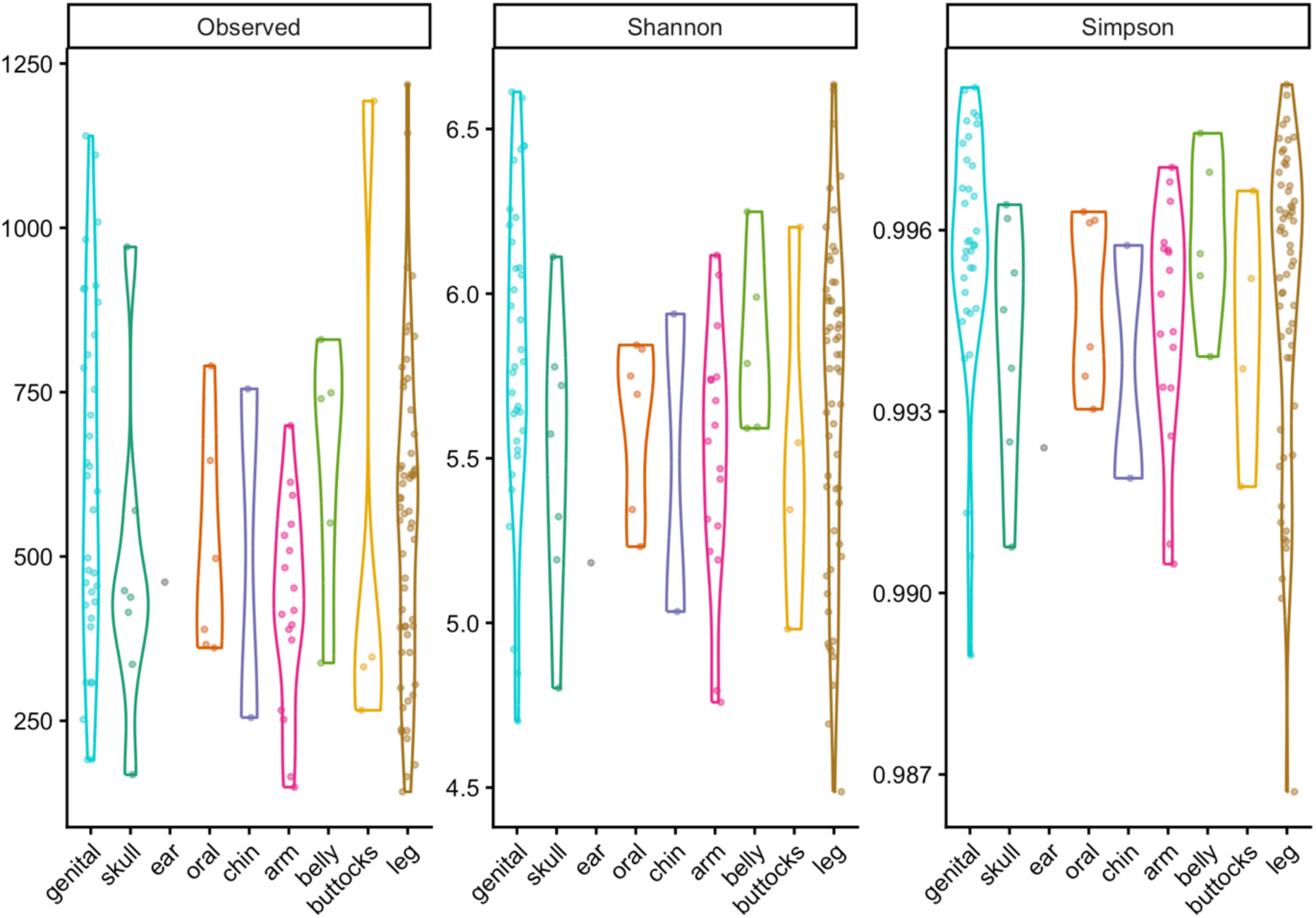
Alpha diversity metrics across different lesion locations. Violin plots of the alpha diversity indices calculated, i.e. observed richness, Shannon and Simpson diversity, calculated for microbial communities sampled from various body sites / individual anatomical locations (genital, arm, belly, buttocks, chin, ear, leg, oral, and skull). Each point represents a skin lesion sample. Statistical assessment of differences in distributions of alpha diversity values was obtained with a Kruskal-Wallis test (*kruskal.test* function in R) a Benjamini-Hochberg correction for multiple testing was applied (*p.adjust* function with method = “BH” in R). While visual differences between individual anatomical locations are apparent, these did not reach statistical significance (adjusted p-values > 0.05).

Differences between genital and non-genital lesions were also investigated with beta diversity analyses using Jaccard and Bray-Curtis distances. As observed in the PCoA plot (**Fig. 4a**, **4b**), the two groups form distinct yet overlapping clusters, where non-genital samples occupy a wider graphical space on the PCoA. Noteworthy, non-genital body site samples do not seem to cluster by skin site (**Fig. 4c**, **4d**). The clustering of genital lesions on the PCoA and the high dispersion observed suggest a wide within lesion group variance, where a large proportion of taxa may not be shared among samples of a group (**Fig. 4** and **5**). As illustrated on **Fig. 5**, the statistical tests for compositional differentiation as well as for dispersion differences were both significant (PERMANOVA adjusted p-values < 0.05; homogeneity of variances adjusted p-values < 0.05). Examination of the distributions of the distances to centroid within lesion groups (**Fig. 5**) highlights both large distances to centroids and wide variance of these distances (especially for non-genital lesions). This suggests some samples harbor taxa that are absent in other samples within their group and / or that some samples have unusual combinations of taxa in comparison to the rest of their group.

**Fig. 4.**
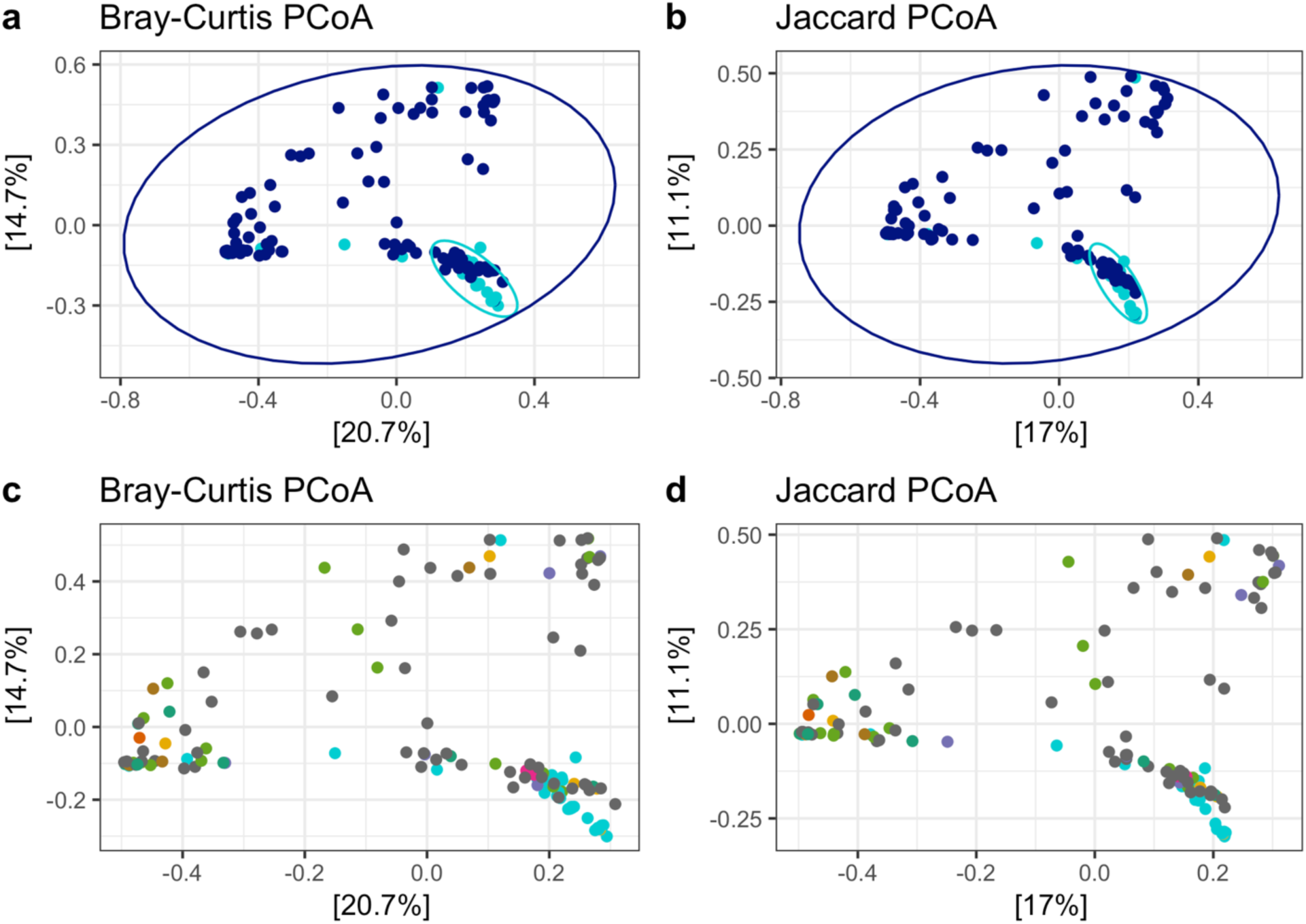
Principal coordinates analysis (PCoA) of beta diversity distances visualising microbial community differences between genital and non-genital lesions. Panels (a) and (b) are coloured by lesion groups (genital vs non-genital as on Fig.2); panels (c) and (d) are coloured by individual anatomical locations (as in Fig. 3). (a) and (c) Bray-Curtis distances, (b) and (d) Jaccard distances; coordinates were obtained with the *ordination* function, *method = PCoA* from the R package **phyloseq**. Ordination plots were produced with the *plot_ordination* function (R package **phyloseq**) and 95% confidence ellipses were drawn with the ggplot2 *stat_ellipse* function. Each point represents the microbial community of a sample.

**Fig. 5.**
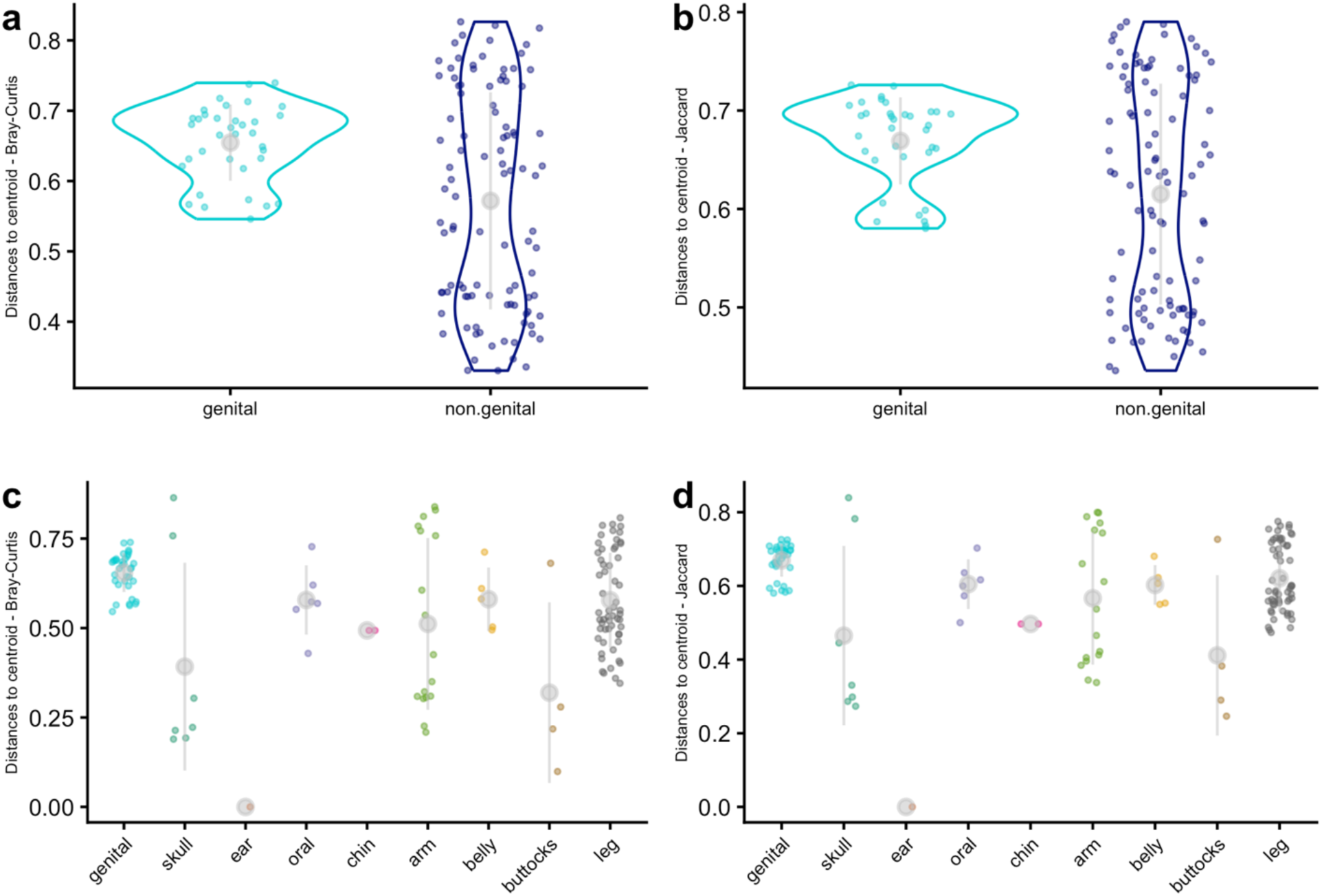
Analysis of homogeneity of group dispersion. Differences in community compositions between genital and non-genital lesions were suggested with PERMANOVA. Therefore, a test of group dispersion was performed (*betadisper* function, R package **vegan**). Distances to group centroids are shown for genital and non-genital lesion groups in panels (a) and (b) – samples colored as in Fig. 2, and for individual anatomical locations in panels (c) and (d) – samples colored as in Fig. 3. Distances to group centroids for Bray–Curtis distances are shown in panels (a) and (c), and for Jaccard distances in panels (b) and (d). Grey circles and vertical bars indicate the mean and standard deviation of the distances to centroid. All analyses rejected the null hypothesis of homogeneity of variances (adjusted p-values < 0.05).

The differences in microbial communities between genital and non-genital skin lesions were further investigated with an analysis of differential abundance measuring the magnitude and direction of change in abundance of genera (**Fig. 6**). This analysis revealed that genital skin lesions tend to be enriched with *Gardenerella*, while non-genital skin lesions tend to be enriched with *Staphylococcus*, *Bradyrhizobium*, *Pelomonas*, *Corynebacterium* and *Streptococcus*. These differentially abundant genera belong to three phyla, i.e. Actinobacteria, Firmicutes and Proteobacteria, that include significant taxa commonly found on the healthy skin. This led to hypothesize that profiles of those taxa commonly found on the healthy skin might differ between genital and non-genital skin lesions. The phyla Bacteroidetes was added to the phyla already mentioned. Within these phyla, specific taxa commonly associated with skin microbiota were sought, such as the families Corynebacteriaceae, Micrococcaceae, and Staphylococcaceae, as well as the genera *Cutibacterium*. The following analyses focused on these taxa to further assess how these ubiquitous communities could vary in skin lesions. Due to the context of the skin lesion samples collected, the phyla Spirochaetes, as well as the genera *Mycobacterium*, and *Haemophilus* were added to the list of microbes of interest.

**Fig. 6.**
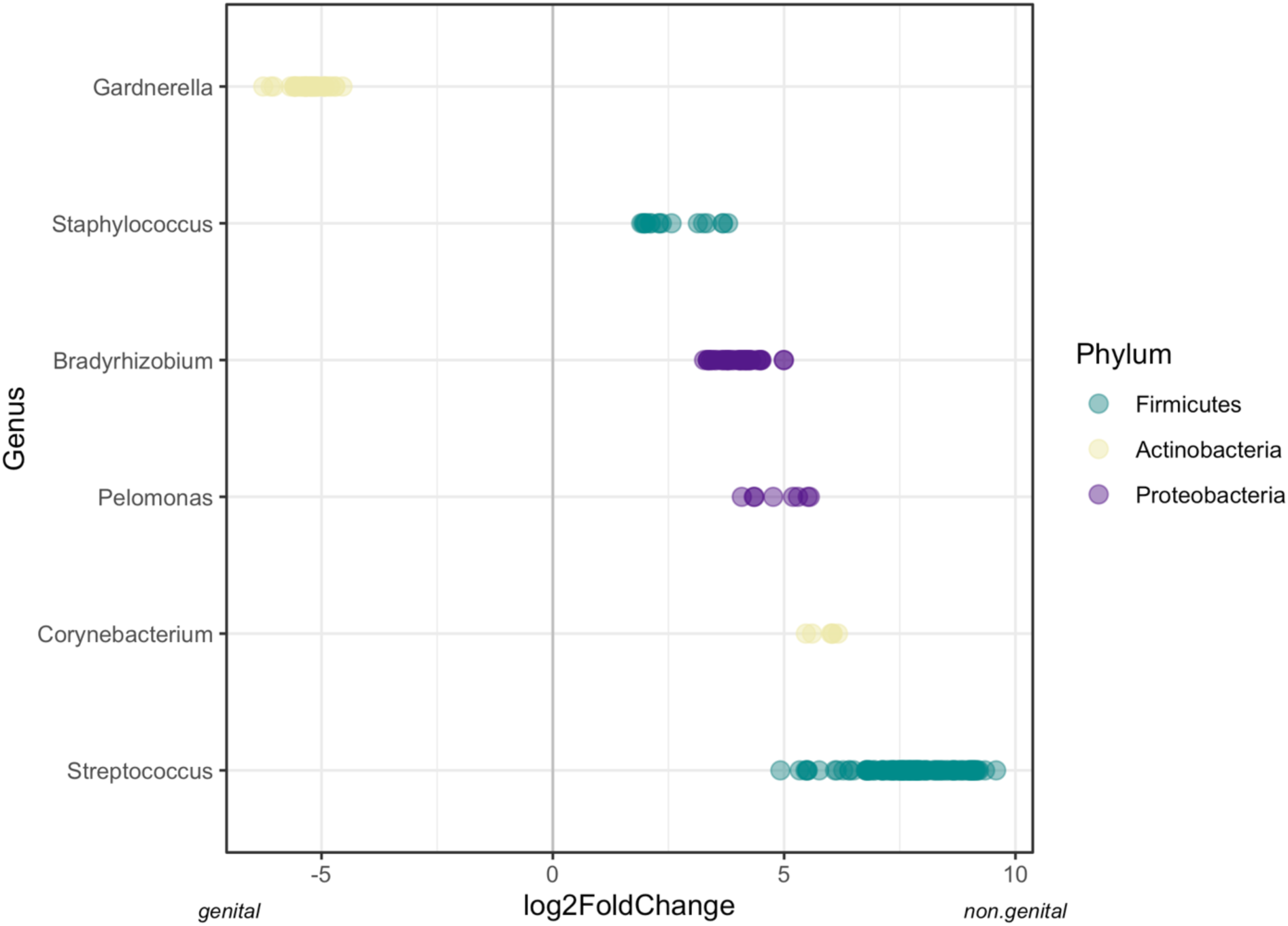
Differentially abundant taxa in genital and non-genital lesions. Points represent different ASVs of the same genus. Plotted log2 fold-change values are DESeq2-estimated log2 fold changes among ASVs, where positive values indicate enrichment in non-genital lesions and negative values enrichment in genital lesions.

### Associations between ubiquitous skin taxa and bacterial diversity in lesions

Considering a skin lesion as a distinct ecological niche to the healthy skin, we investigated how taxa that are usually ubiquitous on the healthy skin could be affected by the lesion microbiota. As a proxy for the lesion microbiota, we used alpha diversity measures, i.e. observed richness, Shannon and Simpson diversity metrics (see Supporting Information **SI1**, **S13-S24 Fig.** and **S Tables 1-5**). The assumption here is that alpha diversity captures some information about the various microbes that co-exist in the skin lesion niche. We investigated whether a relationship between the relative abundance of ubiquitous taxa of the skin and the alpha diversity of skin lesions on genital and non-genital body sites exists. A first linear model was tested to assess the difference in alpha diversity distributions between genital and non-genital skin lesions when considering ubiquitous taxa belonging to Actinobacteria, Bacteroidetes, Firmicutes and Proteobacteria. No significant differences could be evidenced (**SI1**, **S14 Fig.**).

Next, additional linear models were tested to explore the relationship between alpha diversity and the relative abundance of specific taxa of interest (procedure detailed in Supporting Information). Here, the number of reads was used as a proxy for the abundance of a taxon of interest. Within the Actinobacteria phylum, the number of reads from the Corynebacteriaceae and Micrococcaceae families, and from the genera *Cutibacterium* and *Mycobacterium* were counted separately from all other Actinobacteria. Within the Firmicutes phylum, the number of reads belonging to the Staphylococcaceae family was counted separately from other Firmicutes. Within the Proteobacteria phylum, the number of reads of the *Heamophilus* genus was counted separately from other Proteobacteria. The number of reads belonging to the Spirochaetes phyla was also considered. Here, total sum scaling was applied (i.e. relative abundance of the taxa obtained by dividing the number of reads of a taxon of interest per the total number of reads in the corresponding sample). A detailed description of the approach undertaken and of the results obtained is provided in the Supporting Information **SI1** (**S15-S24 Fig.** and **S Tables 1-5**).

The models retained did not suggest body sites captured a significant component of the variance in alpha diversity in relation to the relative abundance of these ubiquitous taxa of the skin (see Supporting Information **SI1**). Nonetheless, these models suggested the relative abundance of Corynebacteriaceae and of Micrococcaceae (both belonging to the Actinobacteria phylum) tend to increase in samples with higher alpha diversity (Shannon and Simpson diversity, **Fig. 7**). A higher diversity (Shannon and Simpson) tends to be observed when both the relative abundance of *Cutibacterium* and those of Micrococcaceae are intermediate (**Fig. 7** and Supporting Information **SI1** – *ActinCutib.prop:ActinMicroc.prop*). However, when the relative abundance of *Cutibacterium* is high, there tends to be relatively fewer Micrococcaceae (**Fig. 7**). A higher diversity tends to be observed when both the number of reads of Staphylococcaceae and those of other Firmicutes are intermediate, when one or the other is very high, diversity (Shannon and Simpson) tends to be low (**Fig. 7** and Supporting Information **SI1** – *FirmiStaphy.prop:OtherFirmicutes.prop).* It should be stressed here that there is no information about what causes one or the other (high alpha diversity on the one hand, intermediate number of reads of Staphylococcaceae and Firmicutes on the other). It nonetheless makes sense that two taxa present in non-negligible amount would lead to a higher value of diversity than the presence of one dominant taxon, unless the one present actually contributes a large number of different species, which is a level of identification that is not available with this 16S data.

**Fig. 7.**
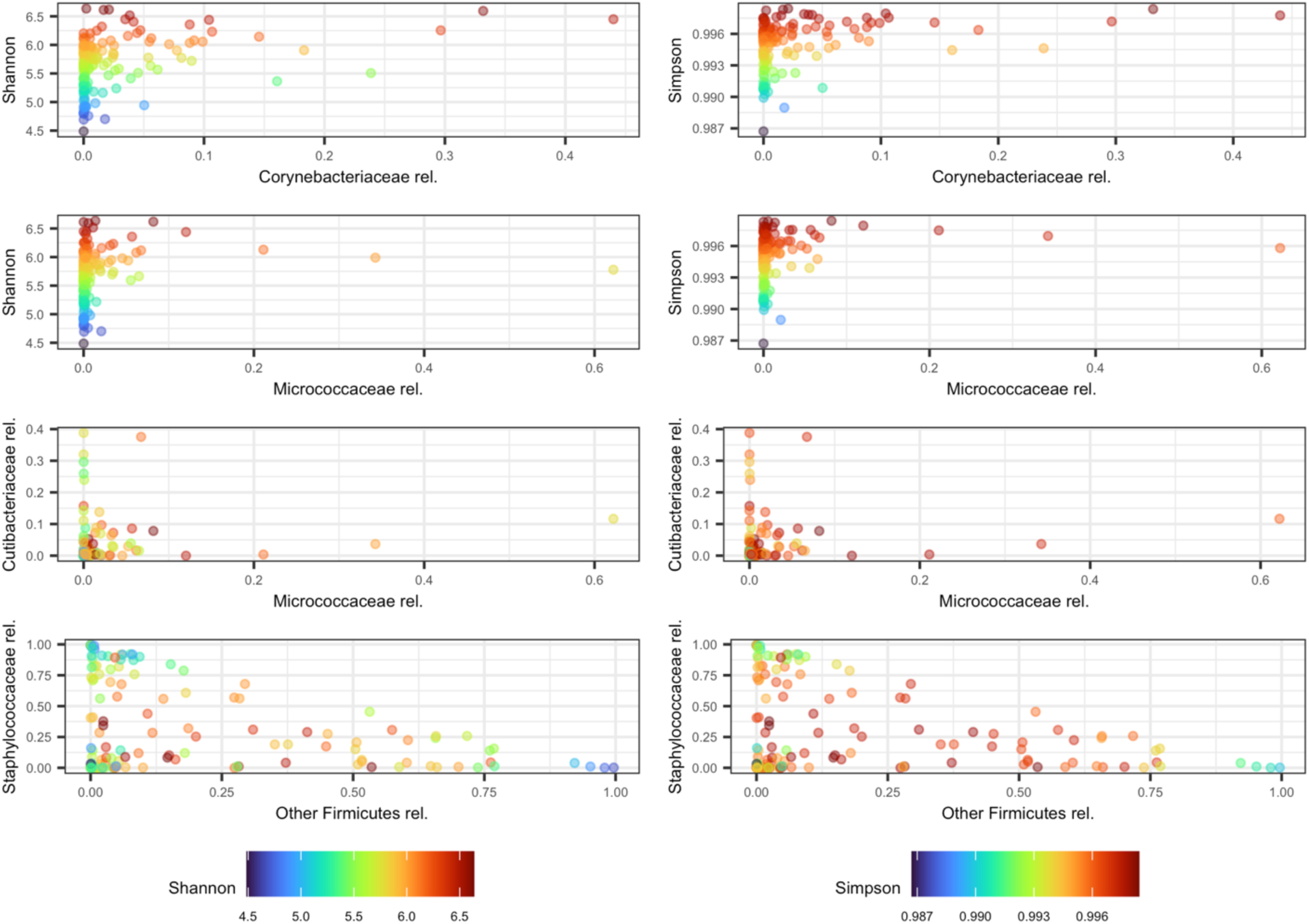
Abundance of ubiquitous taxa of the healthy skin in relation to alpha diversity of skin lesions. The gradient of colour from blue to red show low to high Shannon diversity (left column) and Simpson diversity (right column), respectively. Each point represents a skin lesion sample, colour-coded for the corresponding level of alpha diversity. Only the taxa involved in a significant association with alpha diversity are shown (see Supporting Information). The relative abundances (rel.) of the Corynebacteriaceae and Micrococcaceae families were considered separately from other Actinobacteria present and showed a trend towards a higher relative abundance in samples with higher diversity. Interestingly, higher alpha diversity tends to be observed when the *Cutibacterium* genus (Cutibacteriaceae rel.) and the Micrococcaceae family, both belonging to Actinobacteria, showed intermediate to low relative abundance in skin lesion samples. This is similar what is observed for Firmicutes, where a higher alpha diversity tends to be observed when the relative abundance of the Staphylococcaceae family is intermediate over the relative abundance of other Firmicutes.

## Discussion

This study presents a comprehensive assessment of the microbial composition of genital and non-genital skin lesions in Ghana using 16S rRNA amplicon sequencing. After filtering out lesion samples with fewer reads, 23 phyla, 143 families, and 406 genera were identified in the 135 samples retained, demonstrating that both genital and non-genital skin lesions are colonized by diverse bacterial communities. Although the potential role of host environment and lesion location in shaping microbial community structure has been widely reported for the human skin microbiome (36, 37), this was not evidenced in this study. Previous studies have demonstrated a higher diversity for genital skin lesions could be expected due to the unique environment of the genital region, which is typically moist, lined with mucosal surfaces, and frequently exposed to microbes from the urogenital tract. Alternatively, lesions on the limbs or trunk may not provide the same supportive conditions, as these areas are drier and more exposed to the outside environment, thereby limiting the range of bacteria that can thrive there. However, to our knowledge, the present study is the first to explicitly compare skin microbiomes of genital and non-genital lesions from a region where skin NTDs occur. Albeit the distinct bio-location aspects outlined above and supporting distinct niches, genital and non-genital skin lesions investigated here did not appear to harbor significantly different alpha diversity patterns (Observed richness, Shannon index, and Simpson index).

Beta diversity analyses suggested some differences in microbial community composition between genital and non-genital lesions, as well as differences in dispersion. That is, the microbial communities were not only different on average between lesion types but were also extremely variable within each body site group. Some distances showed more consistency across samples (e.g. distances to centroid of genital lesions consistently high, **Fig. 5**), while others were more heterogeneous (e.g. wide range of low and high distances to centroid of non-genital lesions, **Fig. 5**). This is an important consideration, as it suggests that the observed differences are influenced by both shifts in dominant microbial communities and differences in variability, rather than a single underlying pattern. Interestingly, some of the samples with fewer sequencing reads were still dominated by bacteria like *Treponema*, *Haemophilus*, or *Mycobacterium* microbes known to play a role in skin and soft tissue infections. Even though not all these samples passed our quality threshold for downstream analyses, clinically relevant microbes could be detected, emphasizing 16S sequencing use even when the data yield is low. This highlights its potential value as a diagnostic tool for lesion samples.

Our differential abundance analysis revealed clear differences in the types of bacteria associated with lesion location. Genital (syphilis-like) lesions were enriched with *Gardnerella*, commonly associated with mucosal and urogenital environments. In contrast, non-genital (yaws-like) lesions showed higher abundance of *Staphylococcus* and *Streptococcus*, both of which are well-known skin colonizers and opportunistic pathogens. These organisms may contribute to local inflammation, bacterial load, or delayed healing. In addition, we detected the presence of *Bradyrhizobium*, a soil-associated genus, in non-genital lesions. This likely reflects environmental exposure, particularly in rural settings where contact with soil is common, especially among children.

Members of Actinobacteria, Firmicutes and Proteobacteria are usually part of a stable skin microbiome, as are members of Bacteroidetes. The results of this relative abundance analysis led to hypotheses whether these ubiquitous taxa of the healthy skin were strongly affected by the microbial composition of skin lesions. Indeed, considering these ubiquitous taxa of the skin contribute to the maintenance of a healthy skin barrier, that barrier would be disrupted in a situation when there is a lesion. Using statistical modelling, we examined whether their relative abundance was associated with overall microbial diversity within lesions. Although we did not observe major differences between genital and non-genital lesions when focusing only on these ubiquitous taxa, some interesting patterns emerged. For example, higher microbial diversity tended to be associated with increased relative abundance of Corynebacteriaceae and Micrococcaceae. In addition, diversity appeared to be highest when certain taxa, such as *Cutibacterium* and Micrococcaceae, were present in balanced proportions rather than when one or the other dominating. A similar pattern was observed within the Firmicutes, where an intermediate abundance of Staphylococcaceae and other Firmicute reads associated with higher diversity.

These observations suggest that microbial balance, rather than the presence of a single dominant organism, may be important for maintaining a stable microbial ecosystem. In the context of skin lesions, disruption of this balance could reflect a breakdown of the normal skin barrier and microbial equilibrium. This aligns with broader findings in skin microbiome research, where loss of microbial balance has been linked to disease states and impaired healing (37). Our findings are also consistent with studies of other ulcerative and chronic skin conditions, where polymicrobial communities are commonly observed. For example, in chronic wounds shifts in microbial diversity and the presence of opportunistic pathogens have been associated with delayed healing and disease progression (38, 39). These parallels suggest that microbial interactions within lesions may play an important role beyond the primary causative pathogen. Importantly, these findings raise the possibility that restoring a balanced microbial community could support lesion healing. While antibiotic treatment remains essential for managing infections, future approaches might consider strategies that promote beneficial or commensal microbes as an alternative or a complementary treatment.

It should be stressed that linear modeling in this study was applied to explore associations in a statistically robust manner, as these approaches embed some corrections for multiple testing. There is no information in the models tested allowing to determine the primary cause of the ulcers or the factors responsible for higher or lower microbial diversity of the ulcers. Linear modelling is not used here as a predictive tool and further studies using independent datasets – especially including samples of adjacent healthy skin – will be needed to validate these observations. The approach undertaken here has allowed to formulate novel hypotheses using an original dataset that need to be supported further with distinct sets of skin lesion samples and complementary data collection.

### Perspectives / Further considerations

This study compares microbiome profiles of genital and non-genital skin lesions from Ghana, a country where the skin NTD yaws is endemic. No significant difference in alpha diversity was evidenced between genital and non-genital body sites, while significant differences in beta diversity were observed. Some differences were highlighted for specific combinations of pathogens, that are usually ubiquitous on the skin. This suggests further signs of the disruption of the skin protective barrier should be investigated rather than the presence / absence of one dominant pathogen. In the future, information about the severity of the ulcer (e.g. including depth and width of the wound) may be considered to relate alpha diversity levels (e.g. high or low) to how close to healing the ulcer is. Hence, even though a balanced and healthy microbiome has been associated with more diverse communities (40), this cannot be formally concluded from the data collected here. Future studies must include skin samples from comparable healthy body sites as well as some information about the severity of the wound for assessing the extent of the skin barrier’s disruption.

The absence of significant differences between genital and non-genital body sites (and among non-genital body sites) suggests that either microbial diversity is broadly similar in skin lesion samples across body sites, and / or that some of the sample sizes per body site are too low to reach statistical significance (e.g. one sample on the ear, two samples on the chin). Furthermore, the skin lesion samples were constituted of about 2.5 times more non-genital lesions than genital lesions, where non-genital lesions include various ecological types of sites, for example on the leg, the knee qualifies as a dry site, while the kneepit (behind the knee) qualifies as a moist site, and similarly on the arm, the elbow qualifies as a dry site, while the inside of the elbow qualifies as a moist site. These distinctions may have brought a distinct understanding of the distribution of the communities observed here.

It should be stressed here that 16S data provides some understanding of the bacterial communities present in genital and non-genital skin lesions, but it leaves out fungi, other Eukaryotes, and viruses, which are all important components of the skin microbiome. Indeed, six of these skin lesions have shown positive amplification of the viruses HSV-1 and −2 DNA (26). Along the same lines, the 16S data collected here has allowed limited taxonomy identification (down to the genus level) while species /sub-species identification would provide a clearer understanding of the pathogenic status of the bacteria. For example, *Staphylococcus* (Firmicutes) are commensal bacteria on the healthy skin. However, increased abundance of *S. aureus* combined with a depletion of *S. epidermis* and *Corynebacterium* spp. was more often observed in patients with atopic dermatitis (AD, a chronic inflammatory dermatosis) in comparison to healthy controls (41). A specific identification of the relative abundance of these genera at the species level, would contribute explaining why skin ulcers may not heal easily and how to facilitate their healing. Investigating microbial communities that are most often observed together in ulcers that are not healing easily may also provide clues into identification of alternative treatments to antibiotics. Those alternative treatments include pro/pre-biotics that have been shown to help a more diverse microbial communities to thrive, thereby supporting a balanced and diverse healthy microbiome (40).

## Conclusion

The microbial diversity of genital and non-genital lesions associated to cases of treponematose in Ghana was compared using 16S DNA sequencing. The analyses performed did not evidence specific anatomical locations were consistently dominated by a single bacterial taxon. Interestingly, differences in relative abundance of some taxa ubiquitously found on the healthy skin were observed between genital and non-genital lesions. While no significant association of a specific pathogen to genital or non-genital lesion could be evidenced, our findings present skin lesions as ecological niches containing diverse bacterial communities that may be involved in the disruption of the healthy skin microbiome.

## Acknowledgements

We thank Dr. Nunes for insightful discussions regarding the analysis of this work, Prof. Sanchez-Mazas for providing infrastructural support and research space during this study, Ms. Oksana Kharabora for supporting molecular laboratory activities, and Prof. Arlene Seña for providing support for laboratory training.

## Supporting information

### Supporting information file SI1

S1 Fig. Summary of steps involved in the library preparation for the 16S rRNA gene sequencing.

S2 Fig. The target sequence amplification of the first 90 samples showing the amplifications of samples in first 20 cycles and each subsequent 5 cycles until the total 40 cycles (at the 5th PCR) were reached.

S3 Fig. PCR plate set up showing how the index primer sets were added.

S4 Fig. Amplification of 90 samples after addition of barcodes in five cycle PCR reaction.

S5 Fig. Spreadsheet used to record the fluorescence standard (Zymo Research, USA).

S6 Fig. Spreadsheet used to automatically generate the standard curve equation (Zymo Research, USA).

S7 Fig. The spreadsheets used to record the fluorescence readings of the samples (A), calculated the concentrations of the samples(B) and used to enter the desired concentration of DNA for normalization (C) (Zymo Research, USA).

S8 Fig. The spreadsheets used to automatically calculate the required volume of each sample for normalization (A) and the total volume of pool library as well as the volume of Select-a-Size MagBead buffer required for final library clean-up (B).

S9 Fig. The final average amplicons size (490) in this study (156 samples) after addition of barcodes.

S10 Fig. Rarefaction curve.

S11 Fig. Total number of reads in genital and non-genital skin lesion samples.

S12 Fig. Phylum observed in genital and non-genital skin lesions.

S13 Fig. Phylum abundance in genital and non-genital skin lesions.

S14 Fig. Alpha diversity of genital and non-genital skin lesions, considering ubiquitous taxa of the skin microbiome.

S15 Fig. Diagnostics statistics of the ob.m2.inter.3 model.

S16 Fig. Diagnostics statistics of the model ob.m284.inter.3.

S17 Fig. Diagnostics statistics of the model sh.m2.inter.3.

S18 Fig. Diagnostics statistics of the model sh.m284.inter.3.

S19 Fig. Dot-and-Whisker plot of sh.m284.inter.3 regression model results.

S20 Fig. Relative abundance of taxa of interest and Shannon diversity values in skin lesions.

S21. Fig. Diagnostics statistics of the model si.m2.inter.

S22 Fig. Diagnostics statistics of the model si.m2.inter84.

S23 Fig. Dot-and-Whisker plot of si.m2.inter84 regression model results.

S24. Fig. Relative abundance of taxa of interest and Simpson diversity values in skin lesions.

S Table 1. Multiple and adjusted R squared of some of the models tested with all 135 samples or all samples minus Y009, Y048 and Y051.

S Table 2. Alpha diversity ranges and standard deviation for observed richness, Shannon diversity and Simpson diversity measures in skin lesion samples.

S Table 3. Coefficient estimates, 95% confidence intervals and adjusted p-values of the sh.m284.inter.3 model.

S Table 4. ANOVA comparisons of models si.m1, si.m1v2 and si.m1.inter.2.

S Table 5. Coefficients estimated by the model si.m2.inter84.

### Supporting information file SI2

R script developed and used to analyze the 16S rRNA data.

### Supporting information file SI3

RData file with the original 16S rDNA data and the complete R workspace produced by SI2 and related to this manuscript.

## Data Availability

All relevant data are within the manuscript and its Supporting Information files SI2 SI3.

