## Supplementary information file for "Microbial community profiles of genital and non-genital skin lesions from Ghana"

### 1. 16S ribosomal RNA gene amplification and sequencing - library preparation

Library preparations were performed according to the manufacturer's instructions (Zymo research USA) and it involved five main setups, namely: i. targeted sequence amplification ii. enzymatic clean up iii. barcode addition iv. Library quantification and pooling and v. final library clean-up (**S1 Fig**).

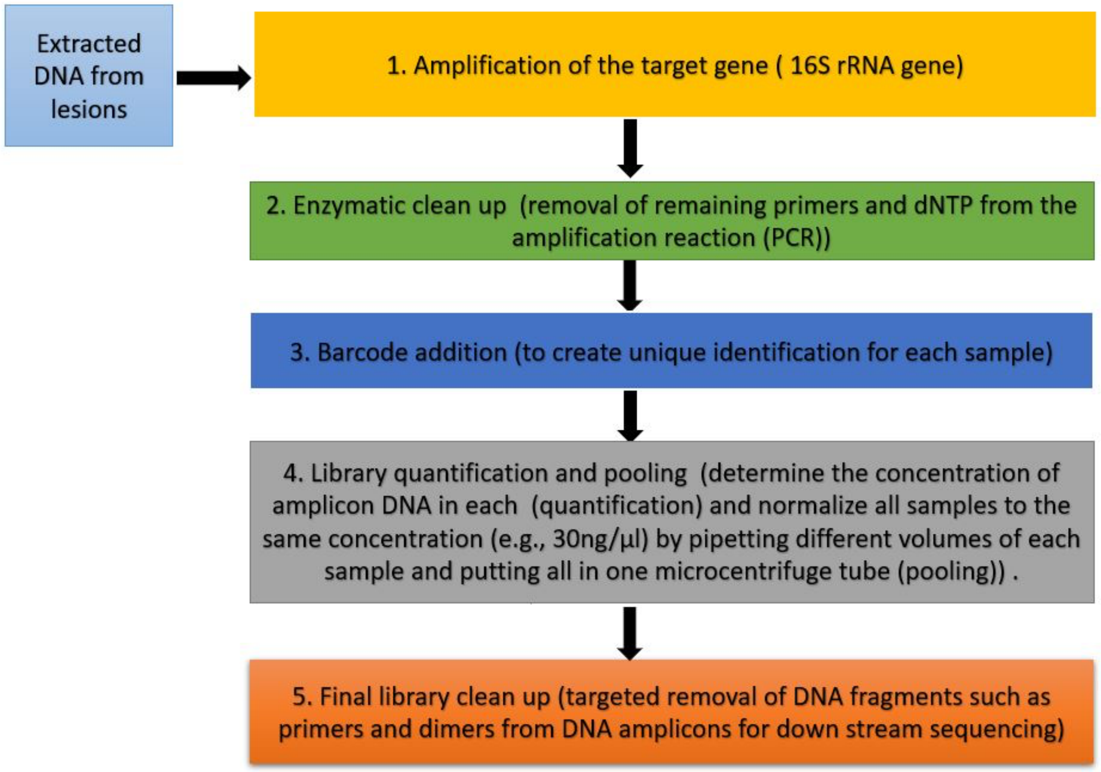


**S1 Fig. Summary of steps involved in the library preparation for the 16S rRNA gene sequencing.**

---

#### 1.1. Targeted sequence amplification

The targeted sequence amplification was performed using 20 µL reaction mixture comprising of 10 µL of Quick-16S qPCR Premix, 4 µL of Quick-16S Primer set V1-V2, 4 µL of ZymoBIOMICS DNase/RNase free water and 2 µL of DNA. The ZymoBIOMICS microbial community standard was used as positive control and DNA free water was used as negative control. The following PCR conditions were used in the BioRad CFX96 real time PCR: initial denaturation at 95 °C for 10 minutes; 20 cycles: 95 °C for 30 seconds, 55 °C for 30 seconds, 72 °C for 3 minutes and plate read. This was followed by 4 °C holding temperature. The fluorescence (from SYBR green fluorophore) from each sample was read after the 20 cycles. Any sample with fluorescence reading above the fluorescent threshold (1200) (**S2 Fig**) of the BioRad

CFX96 real time PCR machine was removed and placed in the same position as it was in the target plate in a new PCR plate labelled collection plate.

The remaining samples that did not pass the fluorescent threshold (1200) were put back into the BioRad CFX96 real time PCR for further 5 cycles using the same conditions. After the 5 cycles those samples that passed the fluorescent threshold (1200) were also removed and place in the same position in collection plate as it was in the target PCR plate. This 5-cycle program was repeated for samples that did not pass the threshold until 40 cycles were reached.

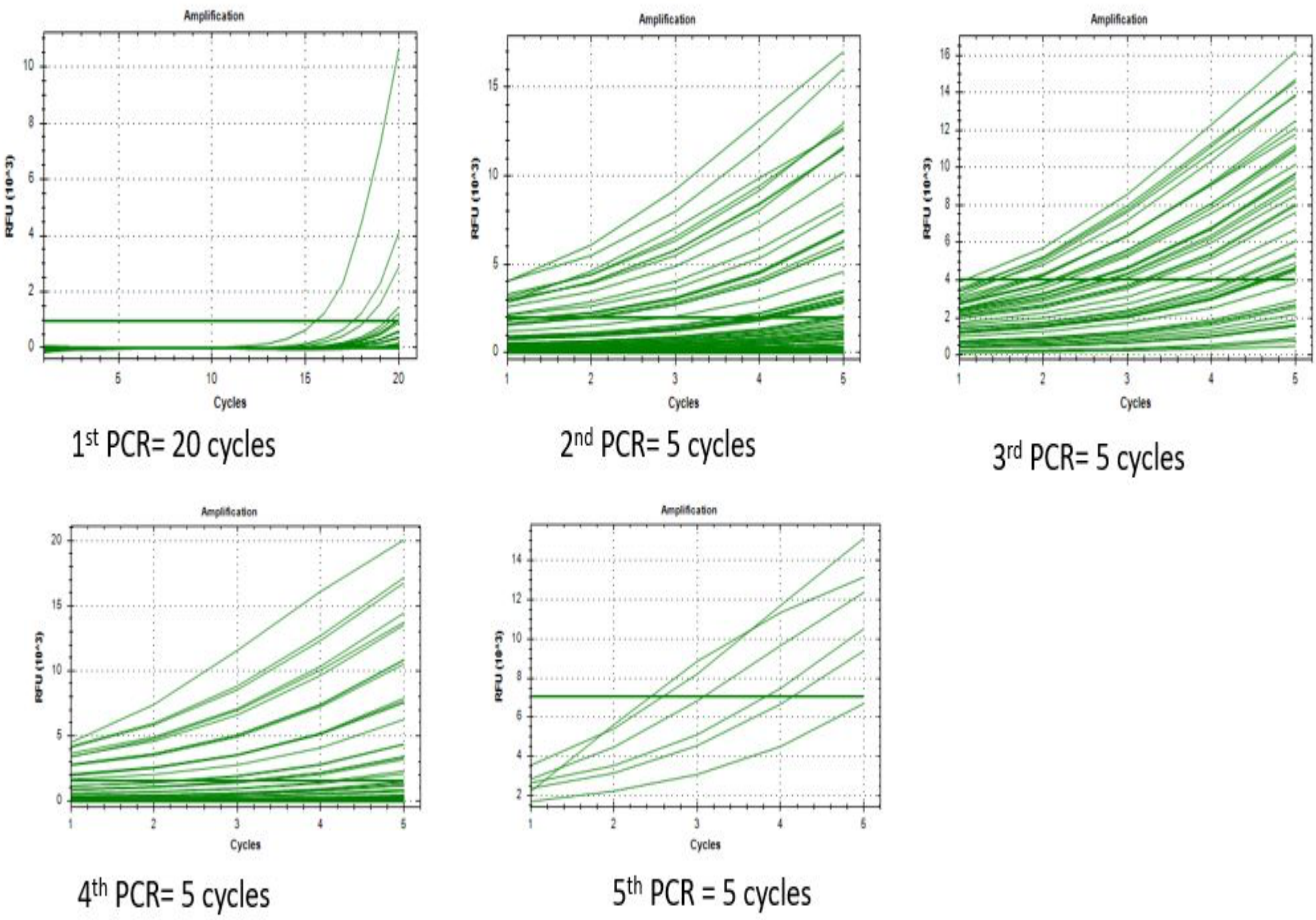


**S2 Fig. The target sequence amplification of the first 90 samples showing the amplifications of samples in first 20 cycles and each subsequent 5 cycles until the total 40 cycles (at the 5$^{th}$ PCR) were reached.**

---

#### 1.2. Enzymatic clean up

The enzymatic cleanup was done by adding 1 µL of reaction clean up solution into each well in the collection plate (containing the amplicons from the targeted sequence amplification step) and the following PCR program were used to run the enzymatic clean up set up in the same real time PCR machine (BioRad CFX96): 37°C for 15mins, 95°C for 10mins and 4°C holding time. After the enzymatic clean-up program, the collection plate was placed on ice for barcode addition.

#### 1.3. Barcode addition

The purpose of the barcoding is to give unique identity to each sample before pooling all samples together for the sequencing. The barcode addition was done with new PCR plate labelled barcoded plate. This was achieved by using 20 µL reaction mixture comprising of 10 µL of Quick-16S qPCR Premix, 4 µL of ZymoBIOMICS DNase/RNase free water, 2 µL of index primer set ZA7xx (ZA701-ZA712) or ZB7xx (ZB701-ZB712), 2 µL of index primer set ZA5xx (ZA501-ZA508) and 2 µL of DNA sample from the collection plate at the end of enzymatic clean program (**S3 Fig**). The following PCR program were used to run the barcode setup in the same real time PCR machine (BioRad CFX96): initial denaturation at 95°C for 10min; 5cycles: 95°C for 30 seconds, 55°C for 30 seconds, 72°C for 3 minutes, plate read and 4°C holding temperature. While the plate was still in the PCR machine, additional program of 72°C for 2mins, plate read and 4°C holding was performed. The fluorescence recordings of each sample well were recorded for library quantification (**S4 Fig**).

Index primer set ZA5xx

Index primer set ZA7xx

| | ZA701 | ZA702 | ZA703 | ZA704 | ZA705 | ZA706 | ZA707 | ZA708 | ZA709 | ZA710 | ZA711 | ZA712 |
|---|---|---|---|---|---|---|---|---|---|---|---|---|
| ZA501 | Y002 | Y023 | Y038 | Y050 | Y080 | Y098 | S006 | S031 | Y008 | Y096 | S022 | S036 |
| ZA502 | Y006 | Y025 | Y039 | Y053 | Y081 | Y101 | S010 | S032 | Y042 | Y099 | S023 | S039 |
| ZA503 | Y007 | Y026 | Y040 | Y056 | Y086 | Y104 | S011 | S035 | Y059 | Y100 | S024 | |
| ZA504 | Y014 | Y029 | Y041 | Y062 | Y089 | Y106 | S012 | S040 | Y064 | Y109 | S026 | |
| ZA505 | Y015 | Y033 | Y044 | Y070 | Y092 | Y108 | S014 | S041 | Y066 | Y110 | S029 | NEG |
| ZA506 | Y019 | Y034 | Y045 | Y073 | Y093 | S002 | S015 | S042 | Y075 | S008 | S030 | |
| ZA507 | Y021 | Y035 | Y046 | Y076 | Y095 | S004 | S016 | S044 | Y079 | S009 | S033 | POS |
| ZA508 | Y022 | Y036 | Y047 | Y077 | Y097 | S005 | S020 | S045 | Y091 | S018 | S034 | POS |

**S3 Fig. PCR plate set up showing how the index primer sets were added**. Each index primer set ZA5xx was added to each well in the row it was positioned (e.g., ZA501 was added to each well from Y002-S036). Each Index primer set ZA7xx was also added to each well in the column it is positioned (e.g., ZA701 was added to each well from Y002-Y022).

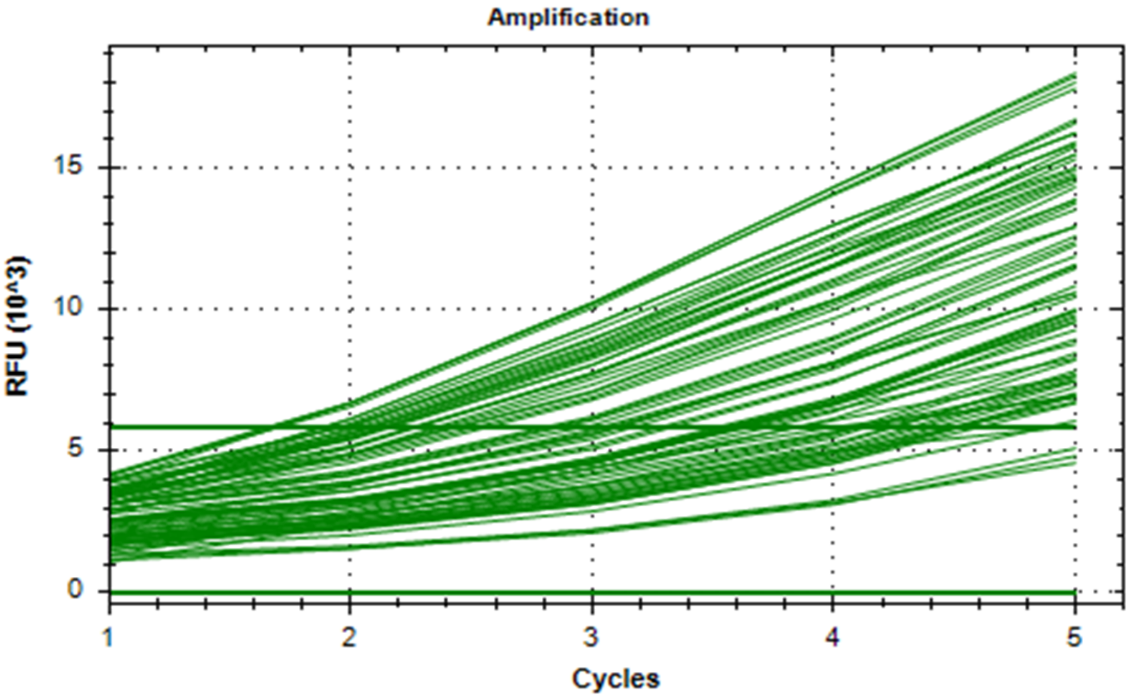


**S4 Fig. Amplification of 90 samples after addition of barcodes in five cycle PCR reaction.**

#### 1.4. Library quantification and pooling

Library quantification and pooling was performed by first recording the fluorescence from the four-fluorescence standard (Zymo research, USA) after running the following program on 20 µL triplicate of each fluorescence standard: 72 °C for 2 mins, plate read and 1 min holding time on same PCR machine. Using the fluorescence reading and the known concentrations (0 ng/ µL, 8ng/ µL, 16ng/ µL and 24ng/ µL) of these standards provided by the manufacturer in addition to the fluorescence readings of each sample obtained from the previous setup (barcoding step), the concentration and volume of each sample needed for the library quantification and pooling were automatically calculated using template spreadsheets provided by the manufacturer (Zymo research, USA) (**S5-S8 Fig**). First, the fluorescence readings of the four standards were recorded on a spreadsheet provided by the manufacture (Zymo research, USA) (**S5 Fig**). The average of the standard fluorescence readings and their standard concentrations were used to create a standard curve equation automatically on a separate spreadsheet (**S6 Fig**). Afterwards, the fluorescence readings of each sample obtained from the previous setup (barcoding step) were recorded on another spreadsheet (**S7A Fig**) and their concentrations were automatically calculated using the standard curve equation on a separate spreadsheet (**S7B Fig**). The concentration (30ng/µL) required for normalization was entered on another spreadsheet (**S7C Fig**) and the volume of each sample for normalization was also automatically calculated on another spreadsheet (**S8A Fig**). After this, the final volume of the pooled library as well as the volume of the select-a-size MagBead buffer needed for the final library clean up were automatically calculated on a different spreadsheet (**S8B Fig**).

| Enter the fluorescence reading (F) values from the "Fluorescence Standards Plate Read" program in the appropriate cells below. | | | | |
|---|---|---|---|---|
| Standard # | 1 | 2 | 3 | 4 |
| F (Replicate 1) | 2355.66 | 7103.76 | 7579.89 | 14345.67 |
| F (Replicate 2) | 2194.21 | 6998.19 | 7688.79 | 14329.21 |
| F (Replicate 3) | 2199.75 | 7063.37 | 7542.62 | 14301.72 |

**S5 Fig. Spreadsheet used to record the fluorescence standard (Zymo Research, USA)**. Each standard fluorescence was prepared and read in triplicate and recorded.

| The average fluorescence reading values will be calculated below and then are used to created a standard curve equation, represented by Values A, B, and C. The R-sqaured value is also calculated and shown. | | | | | |
|---|---|---|---|---|---|
| Average | | 2250 | 7055 | 7604 | 14326 |
| Concentration (ng/µl) | | 0 | 8 | 16 | 24 |
| Value A: | -5.73E-08 | Value B: | 2.96E-03 | Value C: | -6.59301 |
| R-squared: | | | 0.927 | | |

| Array 1 (Used only for R-squared calculation) | |
|---|---|
| 2250 | 0 |
| 7055 | 8 |
| 7604 | 16 |
| 14326 | 24 |

**S6 Fig. Spreadsheet used to automatically generate the standard curve equation (Zymo Research, USA).** The average of each fluorescence standard was automatically calculated by this spreadsheet, and they were used to create a standard curve equatio**n.** The standard equation was used to calculate the unknown DNA concentration for each sample using the fluorescence reading of the sample. The R-squared value indicate how well the standard equation can predict the unknown DNA concentration of the samples. The R-squared value ranges from 0-1 or 0-100%. The higher the value the better. Here 0.927 was obtained.

**A**

Enter the fluorescence reading (F) values from the "Plate Read" program below.

Fluorescence Reading (F)

| | 1 | 2 | 3 | 4 | 5 | 6 | 7 | 8 | 9 | 10 | 11 | 12 |
|---|---|---|---|---|---|---|---|---|---|---|---|---|
| A | 7118.54969 | 11564.83652 | 8247.505865 | 15791.82322 | 10858.45229 | 10029.12305 | 10259.93982 | 12785.88489 | 16493.38266 | 16022.29129 | 10462.96077 | 11067.01662 |
| B | 17536.30736 | 11136.06727 | 8532.127789 | 13804.74618 | 11033.83972 | 16749.77095 | 12848.08707 | 18277.14397 | 15573.97267 | 23793.11024 | 27756.23509 | 14341.94762 |
| C | 20476.93657 | 12208.22938 | 12931.86516 | 21968.7588 | 8680.556543 | 21171.57393 | 18655.01795 | 9112.345911 | 13871.62291 | 14276.02008 | 10962.36211 | 18152.80915 |
| D | 14850.61686 | 6769.288321 | 13619.58922 | 10285.95128 | 18951.02288 | 20373.5911 | 21549.39913 | 11188.57014 | 9118.589053 | 12739.40907 | 27532.71122 | 11338.75801 |
| E | 9067.789386 | 11011.48439 | 22047.19217 | 2904.493609 | 16806.95813 | 18666.88578 | 10983.47512 | 11179.01033 | 15649.26009 | 18888.8108 | 28253.02894 | |
| F | 6580.126967 | 17444.51756 | 24220.90404 | 21732.7173 | 8787.215749 | 11230.7533 | 2400.770806 | 12081.97842 | 6797.861914 | 16195.99004 | 10819.35912 | |
| G | | 9957.64102 | 12638.60722 | 18220.82498 | 18096.62156 | 19245.39874 | 12924.81236 | 13159.26479 | 12233.01271 | 24826.16433 | 14514.25825 | |
| H | | 8861.436954 | 16158.36102 | 15781.95418 | 8294.388936 | 8858.860683 | 8539.46153 | 10773.15634 | 11168.07215 | 17237.43627 | 19006.4803 | 12631.94709 |

**B**

The DNA concentration of each sample will be calculated below in ng/µl.

DNA Concentration (ng/µl)

| | 1 | 2 | 3 | 4 | 5 | 6 | 7 | 8 | 9 | 10 | 11 | 12 |
|---|---|---|---|---|---|---|---|---|---|---|---|---|
| A | 11.60 | 20.02 | 13.96 | 25.93 | 18.84 | 17.37 | 17.79 | 21.94 | 26.71 | 26.19 | 18.15 | 19.19 |
| B | 27.77 | 19.31 | 14.53 | 23.41 | 19.14 | 26.98 | 22.03 | 28.44 | 25.67 | 31.50 | 31.54 | 24.13 |
| C | 30.08 | 21.06 | 22.16 | 30.87 | 14.82 | 30.48 | 28.77 | 15.66 | 23.50 | 24.05 | 19.02 | 28.34 |
| D | 24.79 | 10.85 | 23.15 | 17.83 | 29.00 | 30.02 | 30.68 | 19.40 | 15.67 | 21.87 | 31.59 | 19.65 |
| E | 15.57 | 19.10 | 30.91 | 1.53 | 27.04 | 28.77 | 19.05 | 19.38 | 25.76 | 28.96 | 31.42 | No Sample |
| F | 10.43 | 27.68 | 31.59 | 30.77 | 15.03 | 19.47 | 0.19 | 20.86 | 10.91 | 26.39 | 18.77 | No Sample |
| G | No Sample | 17.24 | 21.72 | 28.40 | 28.29 | 29.23 | 22.15 | 22.49 | 21.09 | 31.68 | 24.36 | No Sample |
| H | No Sample | 15.17 | 26.34 | 25.92 | 14.05 | 15.17 | 14.54 | 18.69 | 19.37 | 27.48 | 29.05 | 21.71 |

**C**

| Enter the amount of product you would like to use (in ng) for normalization below: | |
|---|---|
| Desired amount of product (ng): | 30 |

**S7 Fig. The spreadsheets used to record the fluorescence readings of the samples (A), calculated the concentrations of the samples(B) and used to enter the desired concentration of DNA for normalization (C) (Zymo Research, USA).** The fluorescence readings of the first 90 samples were entered onto the spreadsheet (A) and the concentrations were automatically calculated on the spreadsheet (B)**.** The concentration of DNA (30 ng/µl) for normalisation was also entered on spreadsheet (c).

**A**

Volume to Normalize (µl)

| The volume that should be used for normalization will be calculated below in µl. | | | | | | | | | | | | |
|---|---|---|---|---|---|---|---|---|---|---|---|---|
| | 1 | 2 | 3 | 4 | 5 | 6 | 7 | 8 | 9 | 10 | 11 | 12 |
| A | 2.6 | 1.5 | 2.1 | 1.2 | 1.6 | 1.7 | 1.7 | 1.4 | 1.1 | 1.1 | 1.7 | 1.6 |
| B | 1.1 | 1.6 | 2.1 | 1.3 | 1.6 | 1.1 | 1.4 | 1.1 | 1.2 | 1.0 | 1.0 | 1.2 |
| C | 1.0 | 1.4 | 1.4 | 1.0 | 2.0 | 1.0 | 1.0 | 1.9 | 1.3 | 1.2 | 1.6 | 1.1 |
| D | 1.2 | 2.8 | 1.3 | 1.7 | 1.0 | 1.0 | 1.0 | 1.5 | 1.9 | 1.4 | 0.9 | 1.5 |
| E | 1.9 | 1.6 | 1.0 | 19.6 | 1.1 | 1.0 | 1.6 | 1.5 | 1.2 | 1.0 | 1.0 | No Sample |
| F | 2.9 | 1.1 | 0.9 | 1.0 | 2.0 | 1.5 | 20.0 | 1.4 | 2.7 | 1.1 | 1.6 | No Sample |
| G | No Sample | 1.7 | 1.4 | 1.1 | 1.1 | 1.0 | 1.4 | 1.3 | 1.4 | 0.9 | 1.2 | No Sample |
| H | No Sample | 2.0 | 1.1 | 1.2 | 2.1 | 2.0 | 2.1 | 1.6 | 1.5 | 1.1 | 1.0 | 1.4 |

**Troubleshooting:** Samples highlighted in red do not have enough DNA to meet the desired amount of product. For samples highlighted in red, pool the entire volume (20 µl). Alternatively, decrease the value in Table 5 (cell R28) until there are no remaining highlighted samples.

**B**

Obtain Select-a-Size MagBead Buffer Volume Here

| | | |
|---|---|---|
| Do not edit. | Volume of Pooled Library (µl) | 166.26 |
| | Volume of Select-a-Size MagBead Buffer (µl) | 133.01 |

**S8 Fig. The spreadsheets used to automatically calculate the required volume of each sample for normalization (A) and the total volume of pool library as well as the volume of Select-a-Size MagBead buffer required for final library clean-up (B).** These volumes were automatically generated after entering the normalization volume (30ng/µl) in S11 C Fig. The concentration of the volume highlighted in red was very low so the total volume (20 µl) in the well was used for the normalization.

#### 1.5. Final library clean up

The final library clean-up was performed by adding 265.34 µL (obtained by picking the volume of each of the 156 samlpes required for normalization together) of library pool to 212.28 µL (0.8 × library pool volume) of Select-a-Size MagBead buffer (mixture of 30 µL of Select-a-Size MagBead Concentrate and 1 mL of Select-a-Size MagBead concentrate). This was mixed thoroughly by vortexing and pipetting until it was uniform. The mixture was incubated at 25°C for 5mins. The resulting homogeneous mixture was placed in a magnetic rack and incubated at room temperature until all the beads had separated from the solution. The supernatant was removed and discarded. 200 µl of DNA wash buffer was added whiles the tube containing the beads and pool library were still on the magnetic rack. The supernatant was removed and discarded. All residual buffer was aspirated with the

10µl pipette. The tube containing the pooled library and the beads was then removed from the magnetic rack and incubated for 3mins at room temperature to remove traces of buffer. 20µl of ZymoBIOMICS DNase/RNase free water was added to the beads and mixed thoroughly by pipetting. This was placed back into the magnetic rack and incubated at room temperature until all the magnetic beads were separated completely from the elute. The supernatant was then transferred into a clean tube as the final library.

#### 1.6. Quality check for the final library

The final DNA amplicon size after addition of barcodes is approximately 486 base pairs according to the manufacturer's library preparation instructions (Zymo research, USA). The tape station was used to measure if the correct expected library length of the DNA amplicons were obtained for sequencing (**S9 Fig**). The library was then diluted to a final concentration of 20 µL and was sequenced using the illumina MiSeq 2 x 250 paired end reads sequencing pipeline.

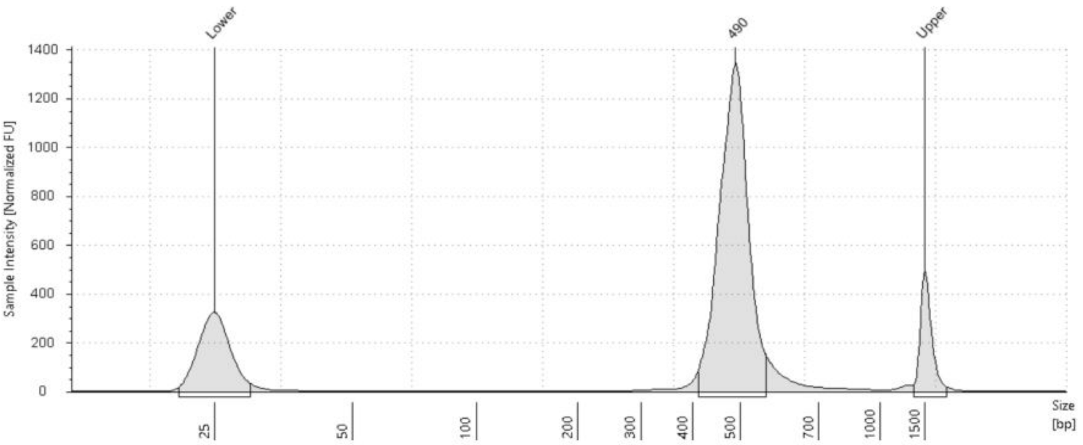


**S9 Fig. The final average amplicons size (490) in this study (156 samples) after addition of barcodes**. The expected amplicon size after addition of barcode according to the manufacturer of the library preparation kits (Zymo Research, USA) is 486.

---

### 2. Analyses of the 16S ribosomal RNA gene data

#### 2.1. Descriptive statistics

The ASV table, the taxonomic table generated by the DADA2 as well as the metadata and the phylogenetic tree were used as input for subsequent analyses in R (version 4.2.2) using the **Phyloseq** (version 1.42.0), **vegan** (version 2.6.4) and **DESeq2** (version 1.38.2) packages. A rarefaction curve was plotted using the **vegan** package to determine if these lesion samples (yaws-like and syphilis-like lesions) were sequenced to a sufficient depth (**S10 Fig.**). Lesion samples with fewer than 1000 reads were not included in downstream analyses (**S11 Fig.**)

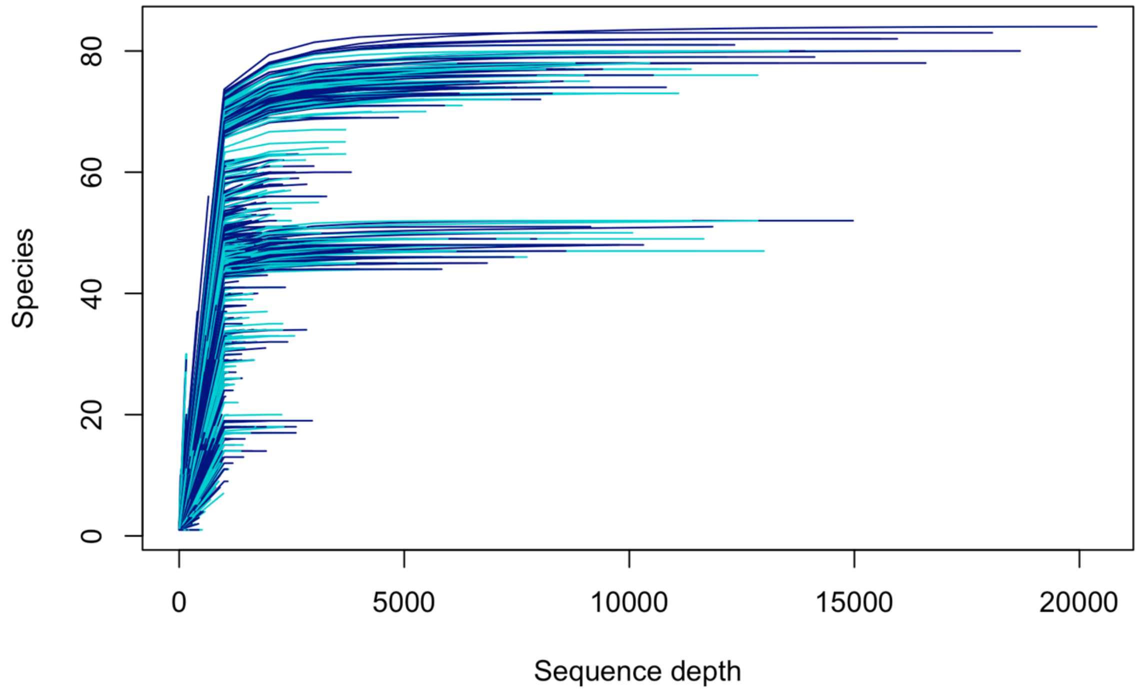


**S10 Fig. Rarefaction curve.**

Colours refer to genital (turquoise) and non-genital (navy ) skin lesion samples.

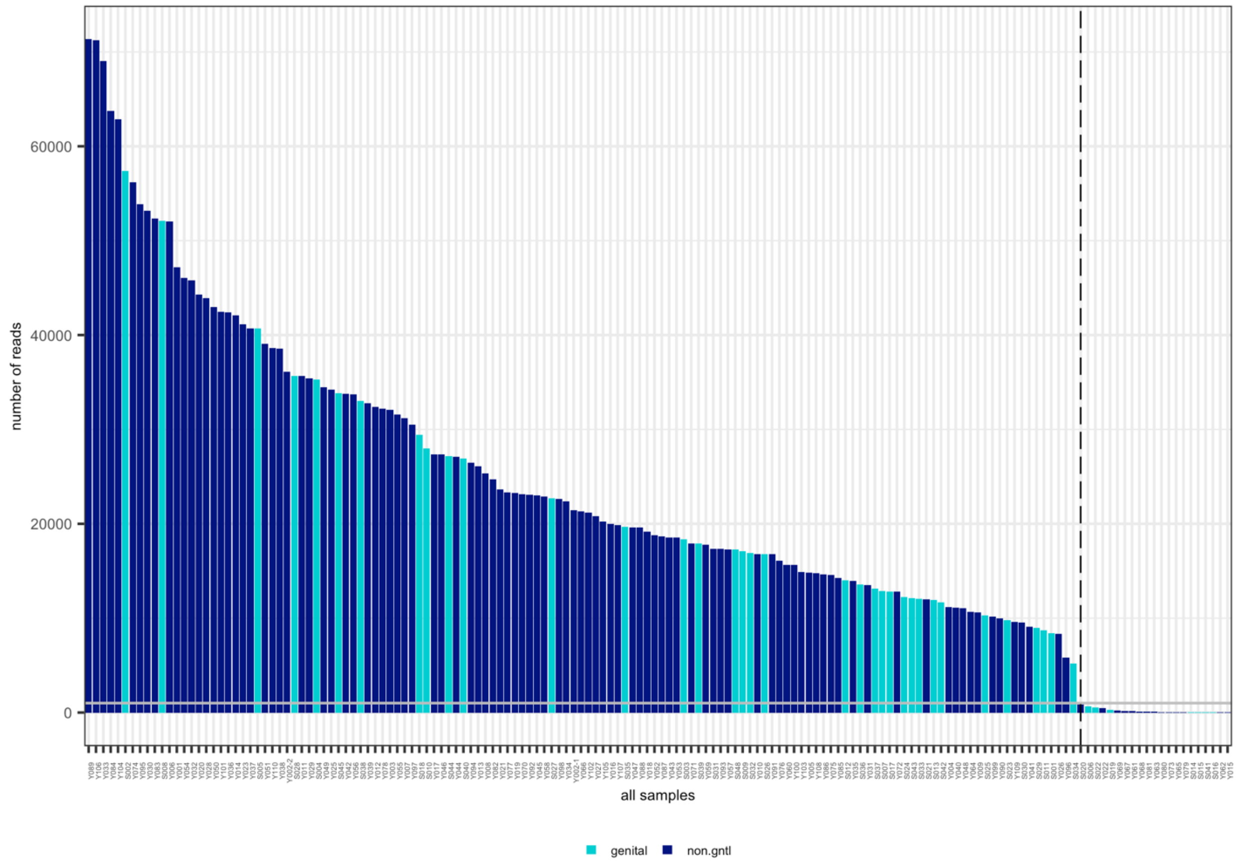


**S11 Fig. Total number of reads in genital and non-genital skin lesion samples.** The y-axis shows the total number of sequencings reads per sample and the bar is colored by sample type (turquoise: genital; navy: non-genital). Samples are ordered from the highest to lowest total read counts. The dashed vertical line indicates a quality control (QC) threshold (1000 reads), below which samples are considered to have insufficient read counts for confident downstream analysis, i.e. 8 genital and 13 non-genital lesions do not pass this threshold.

---

---

Across the 135 samples retained, after removing unidentified phylum sequences, a total of 23 phyla, 143 families and 406 different genera were identified (**Fig. 1**). **Fig. 1** and **S12** and **S13 Fig.** show the same phyla can be observed in both genital and non-genital lesions, although relative abundances appear to differ.

---

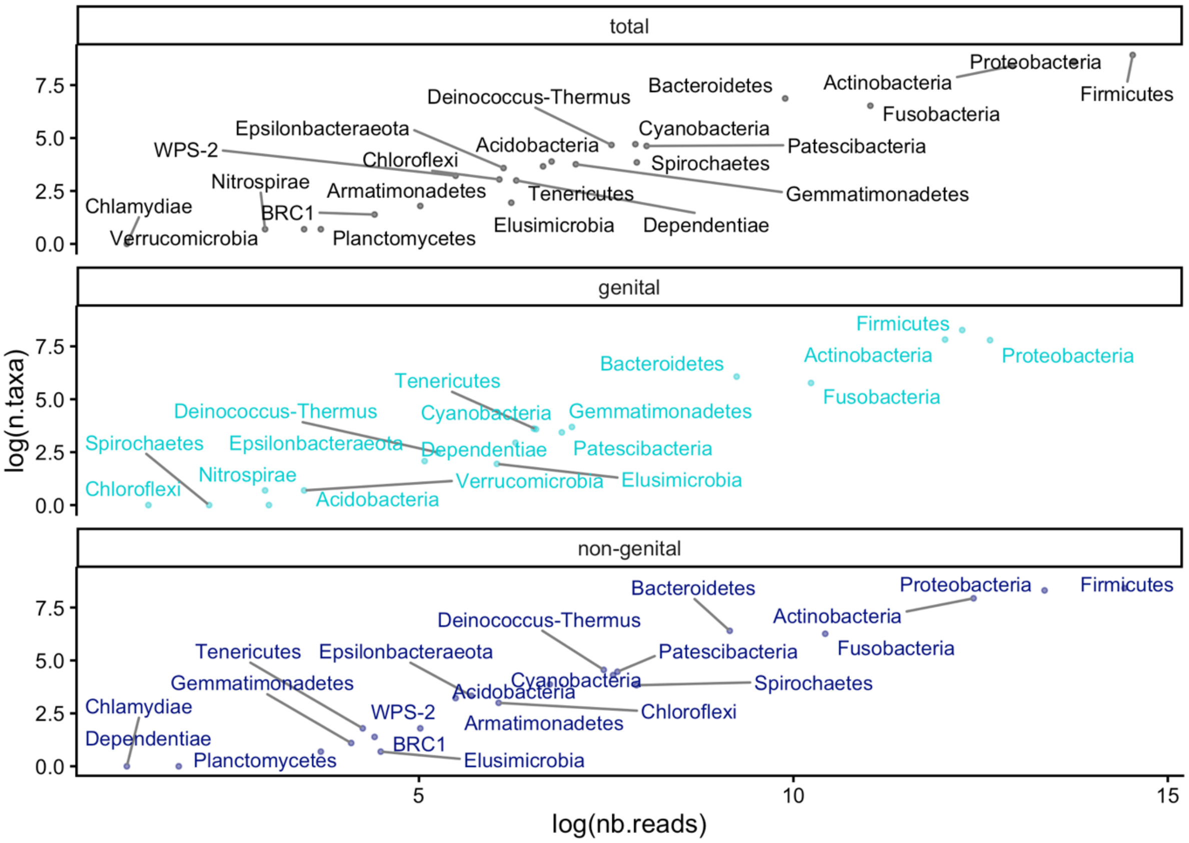


**S12 Fig. Phylum observed in genital and non-genital skin lesions.**

Twenty-three (23) phyla were observed across all 135 skin lesion samples with at least 1000 reads. The x- and y-axes show the natural logarithms of the number of reads and the number of taxa, respectively. (total, top) all 135 lesion samples together; (genital, middle) the 35 genital skin lesion samples with at least 1000 reads; (non-genital, bottom) the 100 non-genital skin lesion samples with at least 1000 reads.

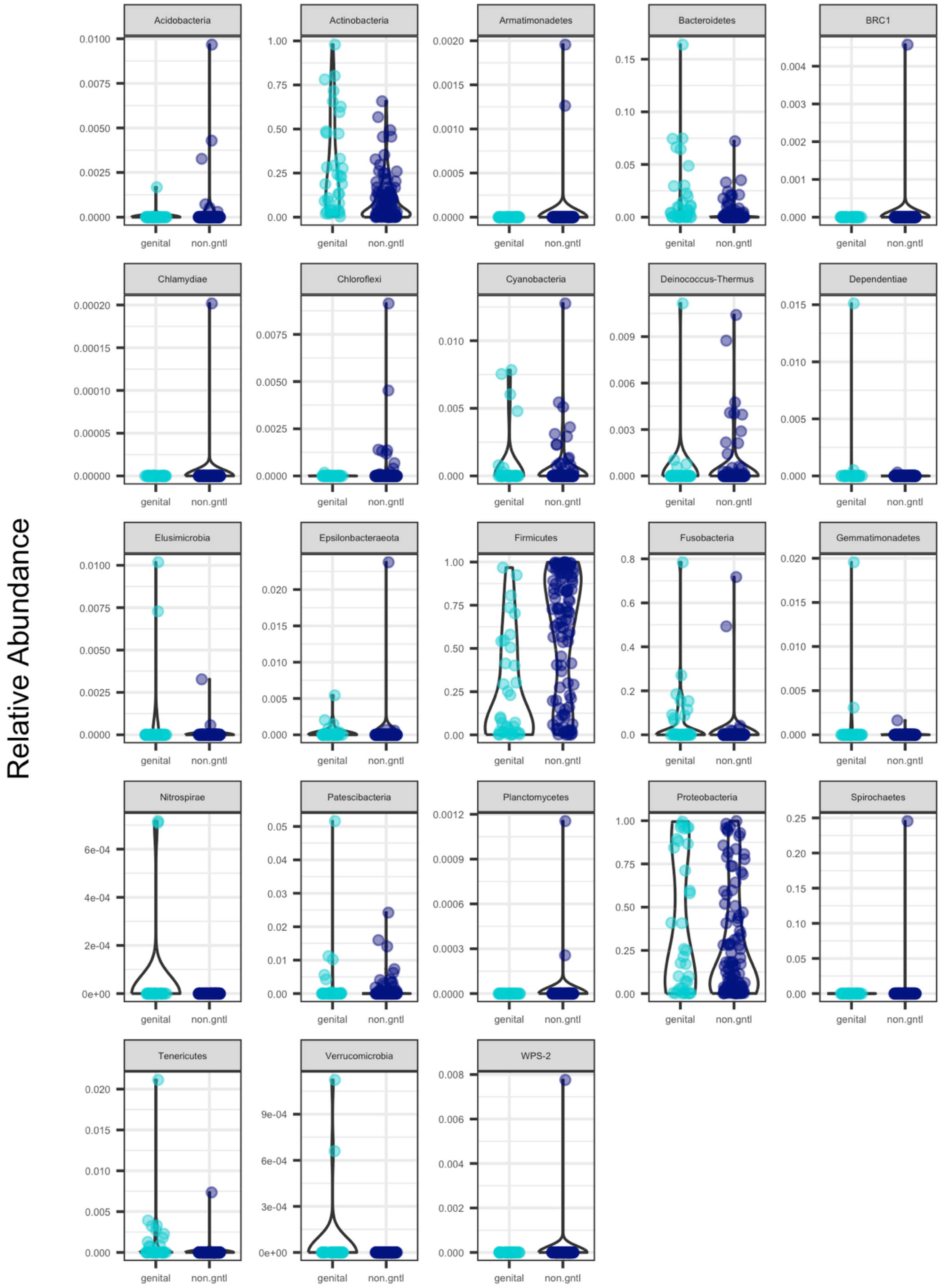


**S13 Fig. Phylum abundance in genital and non-genital skin lesions.**
Relative number of reads (y-axis) of each phylum observed in the 35 genital and the 100 non-genital skin lesion samples (with at least 1000 reads).

---

### 2.2. Analysis of ubiquitous taxa of the skin in genital and non-genital skin lesion samples

The following analyses were performed using the *lm* function to calculate the models, and the function *anova*, to compare them, both functions of base R (R Core Team, 2025). Linear modelling assumptions were verified with the diagnostic plots (function *autoplot*) provided by the R packages **ggfortify** (Horikoshi and Tang, 2018, Tang et al., 2016). Estimates of coefficients of the models retained were plotted with the function *dwplot* from the R package **dotwhisker**.

We investigated how the abundance of some of the ubiquitous taxa usually found in healthy skin microbiomes might be affected in skin lesions. The hypothesis tested here is whether the microbial diversity (i.e. alpha diversity) of skin lesion samples is affected by the relative abundance of specific taxa, known to be ubiquitous in the healthy skin microbiome. This was achieved in two ways. First, we calculated alpha diversity indices (observed richness, Shannon and Simpson diversity measures) only considering those phyla that are known to be ubiquitous taxa of the skin, i.e. Actinobacteria, Bacteroidetes, Firmicutes and Proteobacteria (**S14 Fig.**). A Kruskal-Wallis test was then applied to assess whether alpha diversity measures considering these ubiquitous taxa differed significantly between genital and non-genital skin lesions. None of the three alpha diversity measures suggested there was a difference between genital and non-genital skin lesions based on these ubiquitous phyla (p-values > 0.05). We also applied a linear regression model (function *lm* R software version 4.5.1) to assess if a significant difference in alpha diversity measures considered all together could be identified between genital and non-genital skin lesions. The results of this statistical analysis showed that alpha diversity measures on ubiquitous taxa of the skin could not be used to distinguish the genital/non-genital aspect of the lesions (multiple R-squared: 0.000914, adjusted R-squared: -0.001603, p-value: 0.5471).

---

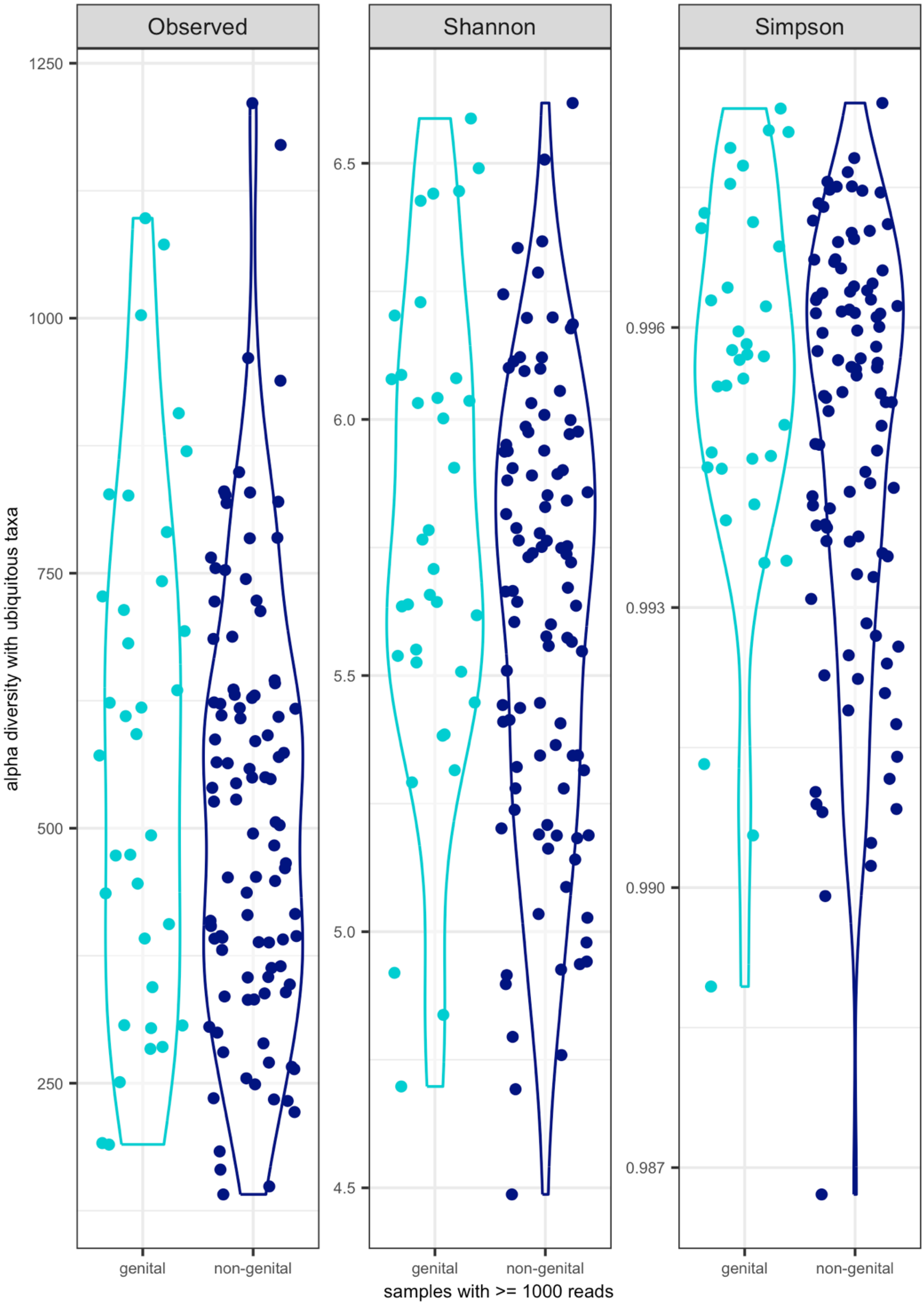
Observed
Shannon
Simpson
alpha diversity with ubiquitous taxa
1250
1000
750
500
250
6.5
6.0
5.5
5.0
4.5
0.996
0.993
0.990
0.987
genital
non-genital
samples with >= 1000 reads

**S14 Fig. Alpha diversity of genital and non-genital skin lesions, considering ubiquitous taxa of the skin microbiome.**
Alpha diversity (observed richness, Shannon and Simpson diversity measures) was obtained considering the ASV numbers belonging to the phyla Actinobacteria, Bacteroidetes, Firmicutes and Proteobacteria. No statistically significant difference between genital and non-genital skin lesions was evidenced (Kruskal-Wallis test and linear regression analysis pvalues > 0.05).

---

Second, we investigated this further by seeking any relationship between alpha diversity and the relative abundance of specific families or genera of interest within those ubiquitous phyla. Within the *Actinobacteria* phylum, we distinguished the number of reads associated to the Corynebacteriaceae and Micrococcaceae families, and down to the *Cutibacterium* and *Mycobacterium* genera from other Actinobacteria. Within the Firmicutes phylum, we distinguished the number of reads associated to the Staphylococcaceae family from other Firmicutes. Within the Proteobacteria phyla, we distinguished the number of reads associated with the *Heamophilus* genus. We further included the number of reads from the Spirochaetes phyla. For each of the 135 skin lesion samples, the relative abundance of these taxa was calculated (number of reads of a taxa of interest / total number of reads obtained in the corresponding skin lesion sample). Alpha diversity values (considering all taxa observed in a sample – and not just those of interest) were obtained for each of the 135 skin lesion samples using the function *estimate_richness* from the R package **phyloseq**. We then applied linear regression analyses to investigate whether there was a relationship between the alpha diversity measures (observed richness, Shannon and Simpson diversity) and the relative abundance of those taxa of interest, also considering the location of the lesion, the qPCR and the DPP test results.

- Observed richness

The following linear model was tested to investigate whether observed richness (total count of unique ASV in a sample, referred to as *observed*) could be affected by the lesion location (*Lesion_location*: genital, arm, belly, buttocks, chin, ear, leg, oral, skull), the *qPCR* result (presence of *Treponema pallidum*, HSV1, HSV2 or *Heamophilus ducreyi* DNA), the *DPP* outcome (positive or negative for ongoing *Treponema* infection) and the relative abundance of any of the taxa of interest:

```
ob.m1: lm(Observed ~ Lesion_location + qPCR + DPP +
        ActinCoryn.prop +
        ActinCutib.prop +
        ActinMicroc.prop +
        ActinMycobact.prop +
        OtherActinobact.prop +
        ProteoHaemoph.prop +
        OtherProteobact.prop +
        FirmiStaphy.prop +
        OtherFirmicutes.prop +
        Bacteroidetes.prop +
        SpirochTrep.prop);
```

where *ActinCoryn.prop* is the relative abundance of Corynebacteriaceae; *ActinCutib.prop* the relative abundance of *Cutibacterium*; *ActinMicroc.prop* the relative abundance of Micrococcaceae, *ActinMycobact.prop* the relative abundance of *Mycobacterium* and *OtherActinobact.prop* the relative abundance of ASV belonging to other Actinobacteria; *ProteoHaemoph.prop* is the relative abundance of *Heamophilus* and *OtherProteobact.prop* the relative abundance of ASV belonging to other Proteobacteria; *FirmiStaphy.prop* is the relative abundance of Staphylococcaceae and *OtherFirmicutes.prop* the relative abundance of ASV belonging to other Firmicutes; *Bacteroidetes.prop* is the relative abundance of *Bacteroidetes*; *SpirochTrep.prop* is the relative abundance of ASV belonging to the Spirochaetes phylum. The function *summary* applied on this model showed it captured some aspects of alpha diversity in the samples with a significant p-value (p-value = 0.013) but relatively low R-squared values (multiple R-squared = 0.286, adjusted R-squared = 0.138) and one term being significantly associated with observed alpha diversity: *ActinCoryn.prop* (p-value = 0.012).

An alternative model was tested with the genital / non-genital (*gntl_nongntl*) location of the lesion rather than the detailed body location argument, as follows:

*ob.m1v2: lm(Observed ~ gntl_nongntl + qPCR + DPP +*
*ActinCoryn.prop +*
*ActinCutib.prop +*
*ActinMicroc.prop +*
*ActinMycobact.prop +*
*OtherActinobact.prop +*
*ProteoHaemoph.prop +*
*OtherProteobact.prop +*
*FirmiStaphy.prop +*
*OtherFirmicutes.prop +*
*Bacteroidetes.prop +*
*SpirochTrep.prop)*

The function summary applied on this model showed it too captured some aspects of alpha diversity in the samples with a significant p-value (p-value = 0.002) but relatively low R-squared values (multiple R-squared = 0.255, adjusted R-squared = 0.154) and two terms being significantly associated with observed alpha diversity: *ActinCoryn.prop* and *Bacteroidetes.prop* (p-values = 0.009 and 0.042, respectively). We next tested a model with interactions where the '*' represents the interactions between terms:

*ob.m1.inter.2: lm(Observed ~ Lesion_location * gntl_nongntl +*
*qPCR + DPP +*
*ActinCoryn.prop **
*ActinCutib.prop **
*ActinMicroc.prop **
*ActinMycobact.prop **
*OtherActinobact.prop +*
*ProteoHaemoph.prop **
*OtherProteobact.prop +*
*FirmiStaphy.prop **
*OtherFirmicutes.prop +*
*Bacteroidetes.prop +*
*SpirochTrep.prop, data = myObserved.df)*

The summary of this model showed it captured some features in the distribution of observed alpha diversity (p-value < 0.001), with relatively low R-squared (multiple R-squared = 0.488, adjusted R-squared = 0.285). Because of the large number of terms estimated by this model, it is not presented here for sake of conciseness. This model was however compared to the other two simpler models with an analysis of variance (ANOVA) using the *anova* function. Both ANOVAs showed the model with interactions was better (p-value of *anova* between *ob.m1* and *ob.m1.inter.2* = 0.003; p-value of *anova* between *ob.m1v2* and *ob.m1.inter.2* = 0.012). We therefore sought to simplify this model by a leave-one-out procedure and comparing the different versions to the complete version (*ob.m1.inter.2*). The terms *Lesion_location*, *gntl_nongntl*, *qPCR* and *DPP* were thereby progressively removed as none of the versions of the model without these seem to affect the original model (all *anova* comparisons showed p-values > 0.05). The model at this stage was expressed as follows:

*ob.m1.inter.2noLqD: lm(Observed ~ ActinCoryn.prop **
*ActinCutib.prop **
*ActinMicroc.prop **
*ActinMycobact.prop **
*OtherActinobact.prop* +
*ProteoHaemoph.prop **
*OtherProteobact.prop* +
*FirmiStaphy.prop **
*OtherFirmicutes.prop* +
*Bacteroidetes.prop* +
*SpirochTrep.prop)*

Formal comparisons of this model to the other two simpler models *ob.m1* and *ob.m1v2* showed the model with interactions (*ob.m1.inter.2noLqD*) was better (*anova* p-values < 0.001 for both comparisons). We consequently tried to remove interactions between three terms and more (using the function *update* as follows):

*ob.m2.inter.3: update(ob.m1.inter.2noLqD, . ~ . -*
*ActinCoryn.prop:ActinCutib.prop:ActinMicroc.prop -*
*ActinCoryn.prop:ActinCutib.prop:ActinMycobact.prop -*
*ActinCoryn.prop:ActinMicroc.prop:ActinMycobact.prop -*
*ActinCutib.prop:ActinMicroc.prop:ActinMycobact.prop -*

*ActinCoryn.prop:ActinCutib.prop:OtherActinobact.prop -*
*ActinCoryn.prop:ActinMicroc.prop:OtherActinobact.prop -*
*ActinCutib.prop:ActinMicroc.prop:OtherActinobact.prop -*
*ActinCoryn.prop:ActinMycobact.prop:OtherActinobact.prop -*
*ActinCutib.prop:ActinMycobact.prop:OtherActinobact.prop -*
*ActinMicroc.prop:ActinMycobact.prop:OtherActinobact.prop -*
*ActinCoryn.prop:ActinCutib.prop:ActinMicroc.prop:ActinMycobact.prop -*
*ActinCoryn.prop:ActinCutib.prop:ActinMicroc.prop:OtherActinobact.prop -*
*ActinCoryn.prop:ActinCutib.prop:ActinMycobact.prop:OtherActinobact.prop -*
*ActinCoryn.prop:ActinMicroc.prop:ActinMycobact.prop:OtherActinobact.prop -*
*ActinCutib.prop:ActinMicroc.prop:ActinMycobact.prop:OtherActinobact.prop -*
*ActinCoryn.prop:ActinCutib.prop:ActinMicroc.prop:ActinMycobact.prop:OtherActinobact.prop)*;

where the ‘:’ represents interactions between terms of the models. We then used the function *anova* which showed this modification did not change the model significantly (p-value > 0.05). However, the diagnostics statistics of this model suggested three samples (Y009, Y048 and Y051) may have a strong influence on the model coefficients (**S15 Fig.**).

---

**S15 Fig . Diagnostics statistics of the *ob.m2.inter.3* model.**

The Residuals vs Fitted plot is a scatter plot of residuals (y-axis) and fitted values (estimated responses, x-axis) that does not show strong non-linearity, unequal error variances, or outliers. The Scale-Location plot displays the fitted values of the regression model on the x-axis and the square root of the standardized residuals on the y-axis. This plot does not indicate that the homoscedasticity assumption is invalid (blue line roughly horizontal) and there is no clear pattern among the residuals. The normal probability plot of the residuals (Normal Q-Q plot) is heavy-tailed, suggesting that the residuals may not be normally distributed. The Cook’s distance plot indicates some specific data points have a strong influence on the model coefficients, as sample numbers 48, 84 and 87 (Y009, Y048 and Y051, respectively) seem to show Cook distances above 0.5.

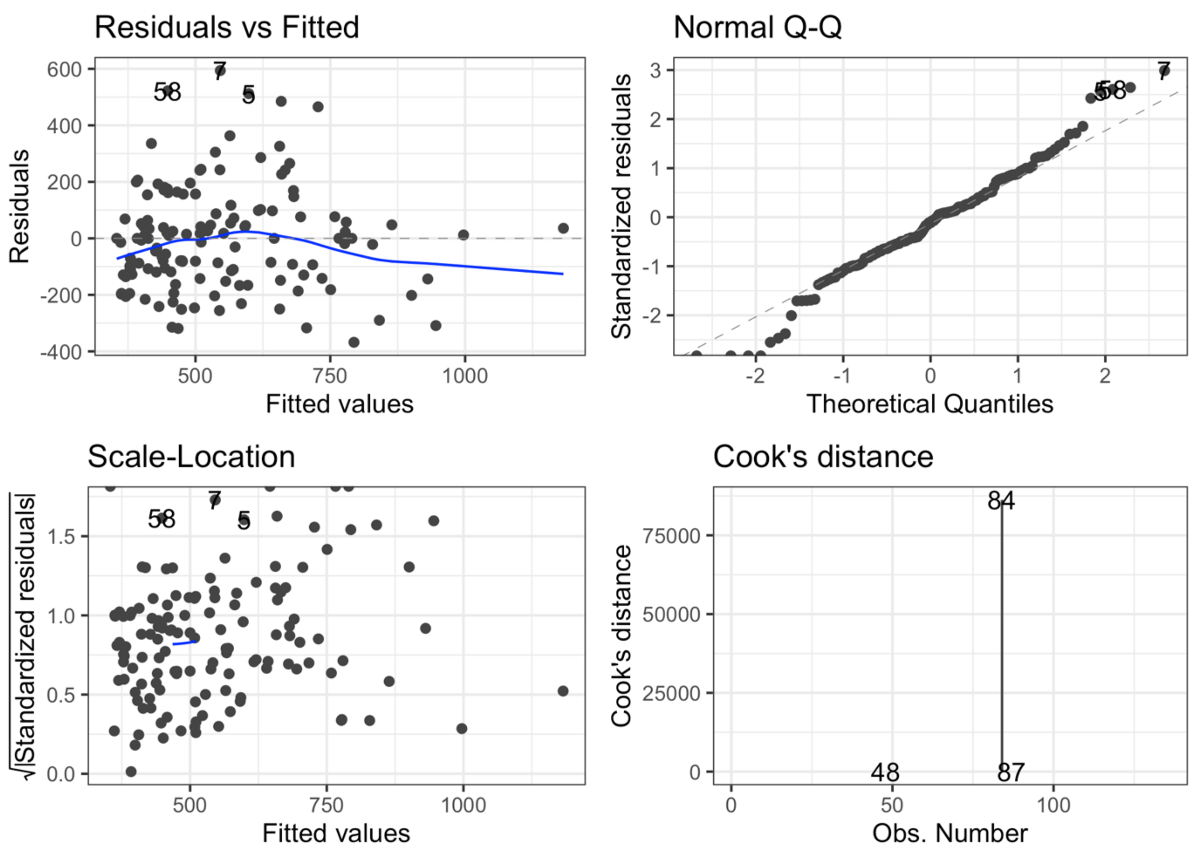


---

These three samples were consequently removed and the models from *ob.m1* to *ob.m2.inter.3* were tested and compared again without these samples (**S Table 1**). They are referred to as *ob.m184* to *ob.m284.inter.3* and include 132 samples. These second set of models were formally compared with *anova* as explained previously. These formal comparisons showed that the model with interactions (*ob.m184.inter.2*) was significantly better than simpler models (*anova* p-values < 0.05). As explained previously, because the model with interactions estimated many terms that were challenging to interpret, this model was simplified throughout a leave-one-out process. Similarly to the approach detailed above, the model with all interactions could be simplified and updated (using the function *update* to remove interactions between three or more terms) without changing the original model significantly (*anova* p-values > 0.05).

---

**S Table 1. Multiple and adjusted R squared of some of the models tested with all 135 samples or all samples minus Y009, Y048 and Y051.**

The models tested with all the samples are shown on the rows showing “135” in the column Samples included. The same models were tested with all the samples except Y009, Y048 and Y051 as those seemed to have a strong influence on the coefficients (132 samples included).

| Model name | Samples included | Multiple R-squared | Adjusted R-squared | p-value |
|---|---|---|---|---|
| ob.m1 | 135 | 0.286 | 0.138 | 0.013 |
| ob.m184 | 132 | 0.296 | 0.146 | 0.011 |
| ob.m1v2 | 135 | 0.255 | 0.154 | 0.002 |
| ob.m184v2 | 132 | 0.262 | 0.159 | 0.002 |
| ob.m1.inter.2 | 135 | 0.488 | 0.285 | < 0.001 |
| ob.m184.inter.2 | 132 | 0.552 | 0.369 | < 0.001 |
| ob.m1.inter.2noLqD | 135 | 0.427 | 0.296 | < 0.001 |
| ob.m184.inter.2noLqD | 132 | 0.507 | 0.391 | < 0.001 |
| ob.m2.inter.3 | 135 | 0.407 | 0.303 | < 0.001 |
| ob.m284.inter.3 | 132 | 0.495 | 0.404 | < 0.001 |

---

The simplified model without the samples (Y009, Y048 and Y051) is referred to as *ob.m184.inter.2noLqD*. The updated model without three or more interactions is referred to as *ob.m284.inter.3*. The *anova* between *ob.m184.inter.2noLqD* and *ob.m284.inter.3* shows the update does not change the model significantly (p-value > 0.05). The diagnostics statistics of this model are shown on **S16 Fig**. They show some of the assumptions of linear modelling are not met and this observed richness may need further transformation to investigate the question of interest.

---

**S16 Fig . Diagnostics statistics of the model *ob.m284.inter.3*.**

The Residuals vs Fitted plot suggests non-linearity, unequal error variances, or outliers. The normal probability plot of the residuals (Normal Q-Q plot) is heavy-tailed, suggesting that the residuals are not normally distributed. The Scale-Location plot does not display clear pattern among the residuals but there is no indication that the homoscedasticity assumption is valid (no blue line). The Cook’s distance plot still indicates that specific data points may have a strong influence on the model coefficients, as distances above 0.5 are observed.

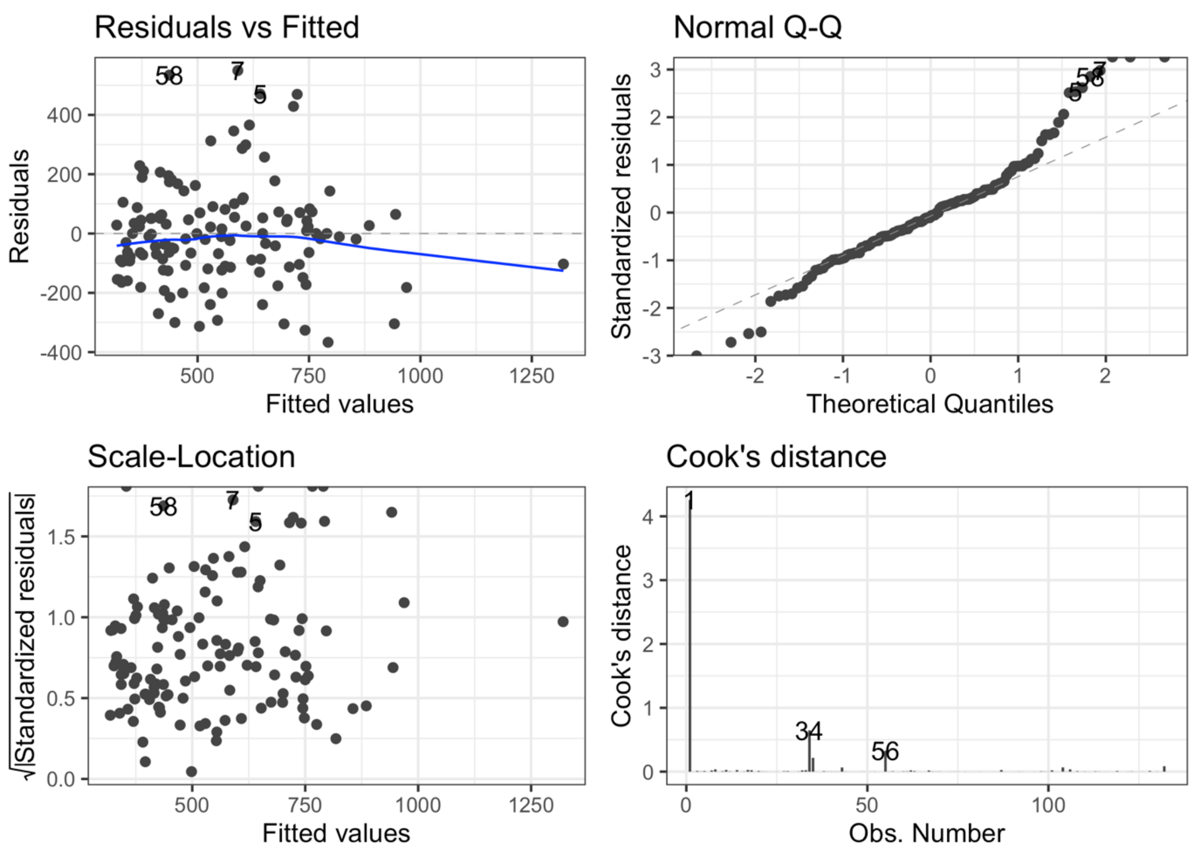


This could also be explained by the wide range of values of observed richness in the 135 samples (from 142 to 1218 ASVs) and a high standard deviation (241.814). Our hypothesis was consequently not pursued further with the observed richness as proxies for skin lesion microbiome diversity values. In comparison, Shannon and Simpson diversity measures harbour narrower ranges and smaller standard deviations, which may make them better suited to investigate our hypothesis (**S Table 2**).

**S Table 2. Alpha diversity ranges and standard deviation for observed richness, Shannon diversity and Simpson diversity measures in skin lesion samples.**

| Alpha diversity | min | max | Standard deviation |
|---|---|---|---|
| Observed | 142 | 1218 | 241.814 |
| Shannon | 4.487 | 6.635 | 0.469 |
| Simpson | 0.987 | 0.998 | 0.002 |

- Shannon diversity index

The same baseline models were tested with Shannon diversity measures. As explained above, the more complex model with interactions captured more of the variation in Shannon diversity measures than simpler models without interactions (both *anova* p-values < 0.01). The complete model can be expressed as follows:

```
sh.m1.inter.2: lm(Shannon ~ Lesion_location * gntl_nongntl +
        qPCR + DPP +
        ActinCoryn.prop *
        ActinCutib.prop *
        ActinMicroc.prop *
        ActinMycobact.prop *
        OtherActinobact.prop +
        ProteoHaemoph.prop *
        OtherProteobact.prop +
        FirmiStaphy.prop *
        OtherFirmicutes.prop +
        Bacteroidetes.prop +
        SpirochTrep.prop)
```

Because this model includes many terms that challenge interpretation, it was simplified in a leave-one-out procedure as explained above. The series of *anova* showed that none of the simplifications changed significantly the model (p-values > 0.05). The following model was consequently favoured:

```
sh.m1.inter.2noLqD: lm(Shannon ~ ActinCoryn.prop *
        ActinCutib.prop *
        ActinMicroc.prop *
        ActinMycobact.prop *
        OtherActinobact.prop +
        ProteoHaemoph.prop *
        OtherProteobact.prop +
        FirmiStaphy.prop *
        OtherFirmicutes.prop +
        Bacteroidetes.prop +
        SpirochTrep.prop)
```

Formal comparisons between *sh.m1.inter.2noLqD* and simpler models without interactions suggested the former should be favoured (*anova* p-values < 0.001). Following a similar reasoning as above, this model was simplified further to remove terms with three interactions or more using the function *update* as follows:

*sh.m2.inter.3: update(sh.m1.inter.2noLqD, . ~ . –*
*ActinCoryn.prop:ActinCutib.prop:ActinMicroc.prop -*
*ActinCoryn.prop:ActinCutib.prop:ActinMycobact.prop -*
*ActinCoryn.prop:ActinMicroc.prop:ActinMycobact.prop -*
*ActinCutib.prop:ActinMicroc.prop:ActinMycobact.prop -*
*ActinCoryn.prop:ActinCutib.prop:OtherActinobact.prop -*
*ActinCoryn.prop:ActinMicroc.prop:OtherActinobact.prop -*
*ActinCutib.prop:ActinMicroc.prop:OtherActinobact.prop -*
*ActinCoryn.prop:ActinMycobact.prop:OtherActinobact.prop -*
*ActinCutib.prop:ActinMycobact.prop:OtherActinobact.prop -*
*ActinMicroc.prop:ActinMycobact.prop:OtherActinobact.prop -*
*ActinCoryn.prop:ActinCutib.prop:ActinMicroc.prop:ActinMycobact.prop -*
*ActinCoryn.prop:ActinCutib.prop:ActinMicroc.prop:OtherActinobact.prop -*
*ActinCoryn.prop:ActinCutib.prop:ActinMycobact.prop:OtherActinobact.prop -*
*ActinCoryn.prop:ActinMicroc.prop:ActinMycobact.prop:OtherActinobact.prop -*
*ActinCutib.prop:ActinMicroc.prop:ActinMycobact.prop:OtherActinobact.prop -*
*ActinCoryn.prop:ActinCutib.prop:ActinMicroc.prop:ActinMycobact.prop:OtherActinobact.prop)*

The *anova* between *sh.m1.inter.2noLqD* and *sh.m2.inter.3* did not show this *update* changed the model significantly (p-value > 0.05). When the diagnostics statistics were investigated, it appeared that the data point 84 may have a strong influence on the coefficient estimates (**S17 Fig.**). We consequently tested the same models but without data point 84 (sample Y048).

---

**S17 Fig . Diagnostics statistics of the model *sh.m2.inter.3*.**

The Residuals vs Fitted plot does not show strong non-linearity, unequal error variances, or outliers. The normal probability plot of the residuals (Normal Q-Q plot) does not suggest the residuals are not normally distributed. The Scale-Location plot does not show a clear pattern among the residuals but it does not indicate that the homoscedasticity assumption is valid as there is no blue line. The Cook's distance plot indicates specific data points may have a strong influence on the model coefficients, as distances above 0.5 are observed (especially for observation 84 – sample Y048).

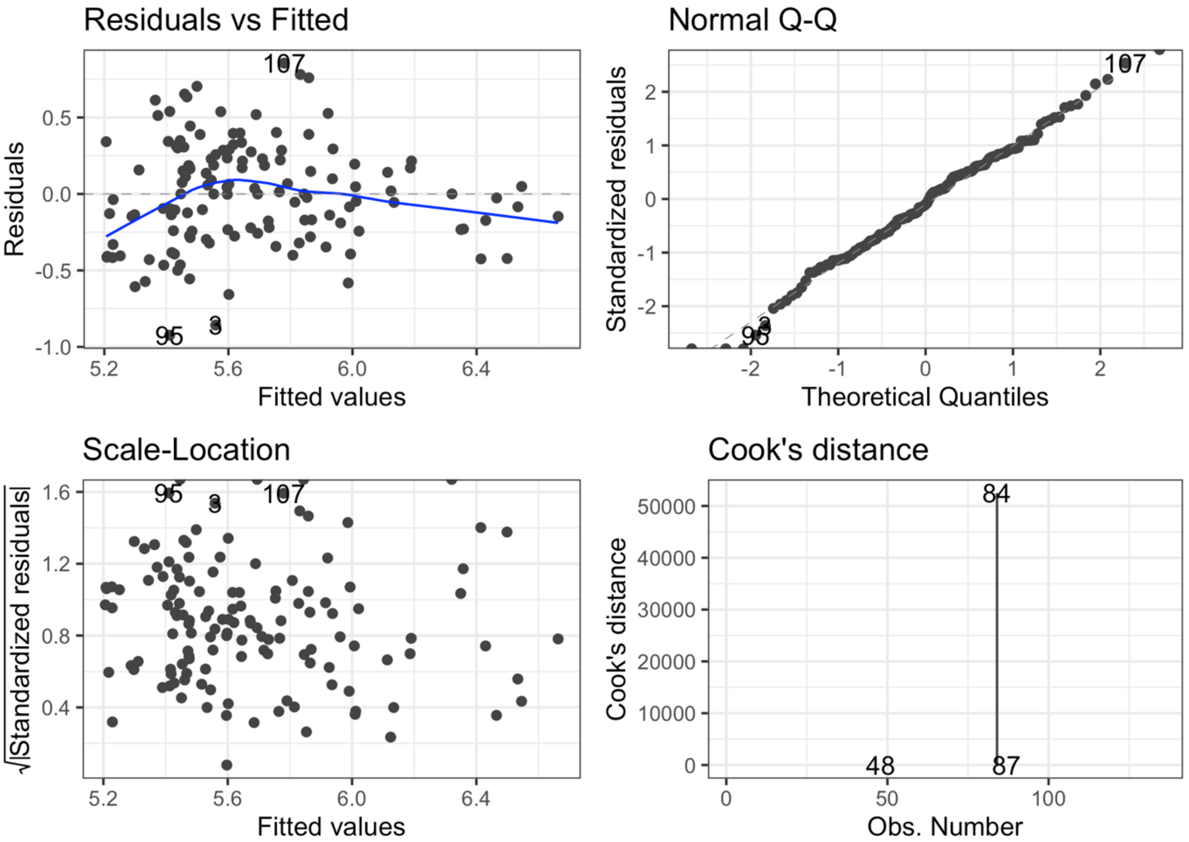


---

We consequently applied the same procedure as explained above, but removing sample Y048 from the data. As detailed above, the following model was the one that was selected after a series of leave-one-out procedure and *anova* comparisons:

*sh.m184.inter.2noLqD: lm(Shannon ~ ActinCoryn.prop **
*ActinCutib.prop **
*ActinMicroc.prop **
*ActinMycobact.prop **
*OtherActinobact.prop +*
*ProteoHaemoph.prop **
*OtherProteobact.prop +*
*FirmiStaphy.prop **
*OtherFirmicutes.prop +*

*Bacteroidetes.prop +*
*SpirochTrep.prop)*

This model was further simplified to remove terms with three or more interactions as follows:

*sh.m284.inter.3: update(sh.m184.inter.2noLqD, . ~ . –*
*ActinCoryn.prop:ActinCutib.prop:ActinMicroc.prop -*
*ActinCoryn.prop:ActinCutib.prop:ActinMycobact.prop -*
*ActinCoryn.prop:ActinMicroc.prop:ActinMycobact.prop -*
*ActinCutib.prop:ActinMicroc.prop:ActinMycobact.prop -*
*ActinCoryn.prop:ActinCutib.prop:OtherActinobact.prop -*
*ActinCoryn.prop:ActinMicroc.prop:OtherActinobact.prop -*
*ActinCutib.prop:ActinMicroc.prop:OtherActinobact.prop -*
*ActinCoryn.prop:ActinMycobact.prop:OtherActinobact.prop -*
*ActinCutib.prop:ActinMycobact.prop:OtherActinobact.prop -*
*ActinMicroc.prop:ActinMycobact.prop:OtherActinobact.prop -*
*ActinCoryn.prop:ActinCutib.prop:ActinMicroc.prop:ActinMycobact.prop -*
*ActinCoryn.prop:ActinCutib.prop:ActinMicroc.prop:OtherActinobact.prop -*
*ActinCoryn.prop:ActinCutib.prop:ActinMycobact.prop:OtherActinobact.prop -*
*ActinCoryn.prop:ActinMicroc.prop:ActinMycobact.prop:OtherActinobact.prop -*
*ActinCutib.prop:ActinMicroc.prop:ActinMycobact.prop:OtherActinobact.prop -*
*ActinCoryn.prop:ActinCutib.prop:ActinMicroc.prop:ActinMycobact.prop:OtherActinobact.prop)*

The *anova* comparison did not suggest this update significantly changed the model. The diagnostics statistics did not show any significant deviation from linear modelling assumptions (**S18 Fig.**).

---

**S18 Fig . Diagnostics statistics of the model *sh.m284.inter.3.***

The Residuals vs Fitted plot does not show strong non-linearity, unequal error variances, or outliers. The normal probability plot of the residuals (Normal Q-Q plot) does not suggest that the residuals are not normally distributed. The Scale-Location plot does not indicate that the homoscedasticity assumption is invalid as there is no clear pattern among the residuals. The Cook's distance plot does not indicate that any specific data points have a strong influence on the model coefficients, as no distances above 0.5 are observed.

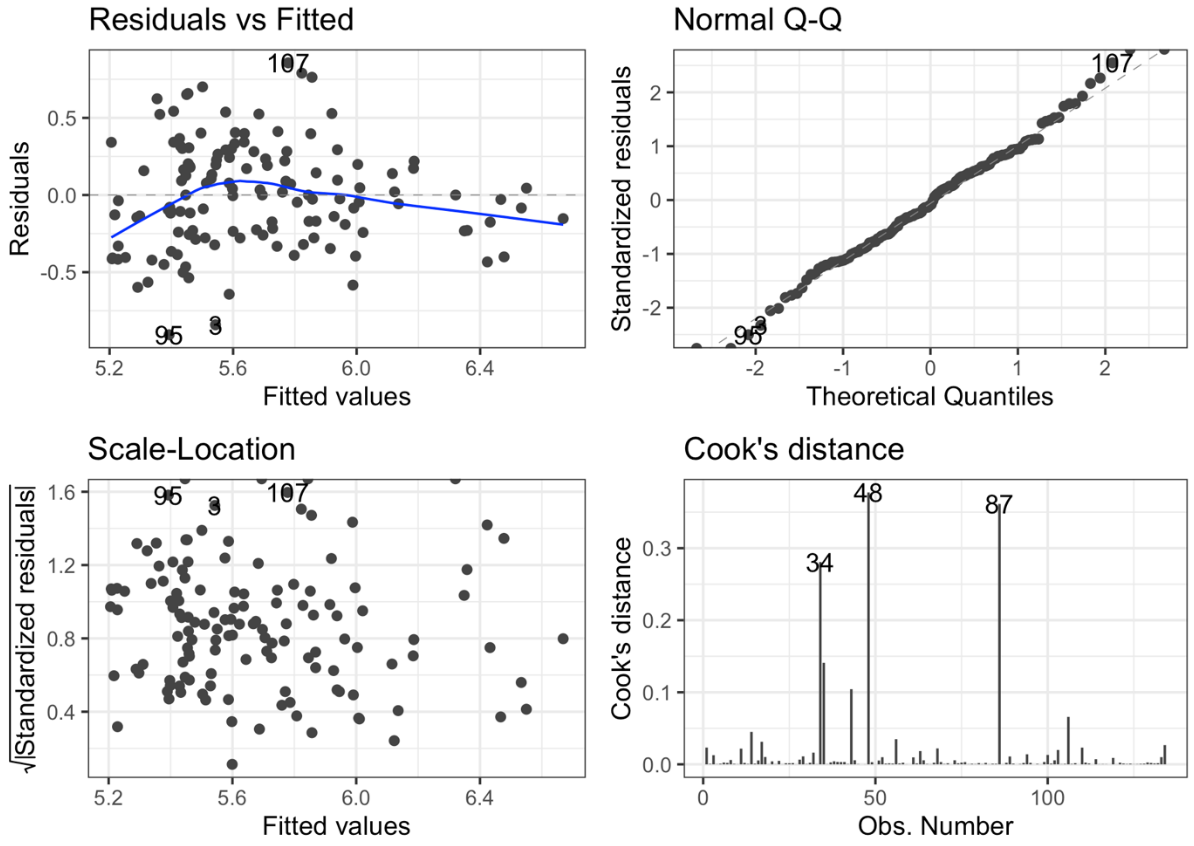


---

The group comparisons of all the groups provided by model *sh.m284.inter.3* are presented in **S Table 3**. The coefficients are presented and discussed below and in the main text.

---

**S Table 3. Coefficient estimates, 95% confidence intervals and adjusted p-values of the *sh.m284.inter.3* model.**

Coefficients in bold are those that are significantly different from 0. Three coefficients could not defined because of singularities. The values provided in the column below were obtained from the *summary* of the model. Residuals: minimum (Min) and maximum (Max), and first (1Q), second (Median) and third (3Q) quantiles of the model's residuals. Estimate: estimate of the coefficient; Std. Error: residual standard error divided by the square root of the sum of the square of that particular x variable; t value: Estimate divided by Std. Error; Pr(>|t|): t value from T distribution table with the given degrees of freedom; Signif. codes: level of statistical significance, 0 '***'

0.001 ‘**’ 0.01 ‘*’ 0.05 ‘.’. For coefficients with interactions (“:”), a positive estimate implies a positive effect (higher mean) of Shannon diversity on the variable to the left, and a negative estimate implies a negative effect (smaller mean) of Shannon diversity on the variable to the left.

| **Residuals** | **Min** | **1Q** | **Median** | **3Q** | **Max** |
|---|---|---|---|---|---|
| | -0.9057 | -2.39E-01 | -2.30E-03 | 0.2256 | 0.8569 |
| **Coefficients** | **Estimate** | **Std. Error** | **t value** | **Pr(>\|t\|)** | **Signif. codes** |
| **(Intercept)** | **5.67E+00** | **3.65E-01** | **15.557** | **< 2e-16** | *** |
| **ActinCoryn.prop** | **1.80E+00** | **7.89E-01** | **2.287** | **0.024061** | * |
| ActinCutib.prop | 1.06E+00 | 7.42E-01 | 1.426 | 0.156697 | |
| **ActinMicroc.prop** | **4.53E+00** | **1.68E+00** | **2.69** | **0.008217** | ** |
| ActinMycobact.prop | 1.35E+03 | 1.26E+03 | 1.068 | 0.287743 | |
| OtherActinobact.prop | -2.40E-02 | 4.52E-01 | -0.053 | 0.957687 | |
| ProteoHaemoph.prop | NA | NA | NA | NA | |
| OtherProteobact.prop | -2.74E-01 | 3.70E-01 | -0.741 | 0.460209 | |
| FirmiStaphy.prop | -4.67E-01 | 3.75E-01 | -1.246 | 0.215421 | |
| OtherFirmicutes.prop | -3.89E-01 | 3.91E-01 | -0.996 | 0.32155 | |
| **Bacteroidetes.prop** | **4.48E+00** | **2.23E+00** | **2.014** | **0.046417** | * |
| SpirochTrep.prop | 1.59E+03 | 1.19E+03 | 1.336 | 0.184184 | |
| ActinCoryn.prop:ActinCutib.prop | -1.17E+01 | 1.68E+01 | -0.697 | 0.487186 | |
| ActinCoryn.prop:ActinMicroc.prop | 2.46E+01 | 3.87E+01 | 0.636 | 0.526185 | |
| **ActinCutib.prop:ActinMicroc.prop** | **-3.49E+01** | **1.08E+01** | **-3.237** | **0.001587** | ** |
| ActinCoryn.prop:ActinMycobact.prop | -3.83E+04 | 4.57E+04 | -0.838 | 0.403694 | |
| ActinCutib.prop:ActinMycobact.prop | -2.09E+04 | 1.93E+04 | -1.082 | 0.281526 | |
| ActinMicroc.prop:ActinMycobact.prop | 2.75E+03 | 9.52E+03 | 0.289 | 0.772794 | |
| ActinCoryn.prop:OtherActinobact.prop | 1.09E+01 | 1.04E+01 | 1.049 | 0.296607 | |
| **ActinCutib.prop:OtherActinobact.prop** | **4.16E+01** | **1.68E+01** | **2.476** | **0.014767** | * |
| ActinMicroc.prop:OtherActinobact.prop | -3.46E+01 | 4.18E+01 | -0.827 | 0.409827 | |
| ActinMycobact.prop:OtherActinobact.prop | NA | NA | NA | NA | |
| ProteoHaemoph.prop:OtherProteobact.prop | NA | NA | NA | NA | |
| **FirmiStaphy.prop:OtherFirmicutes.prop** | **3.09E+00** | **7.81E-01** | **3.961** | **0.000131** | *** |

Residual standard error: 0.3703 on 113 degrees of freedom

Multiple R-squared: 0.4731, Adjusted R-squared: 0.3799

F-statistic: 5.074 on 20 and 113 DF, p-value: 9.293e-09

The coefficients estimated by this model are presented in **S19 Fig**. These coefficients can be better appreciated when placed in the context of the underlying diversity (**S20 Fig.**).

**S19 Fig . Dot-and-Whisker plot of *sh.m284.inter.3* regression model results.** The whiskers show the 95% confidence interval of the coefficient estimates. The x-axis is scaled to provide an optimal overview of the statistically significant (black) and not statistically significant (grey) coefficients. The dashed vertical line shows no effect on Shannon diversity.

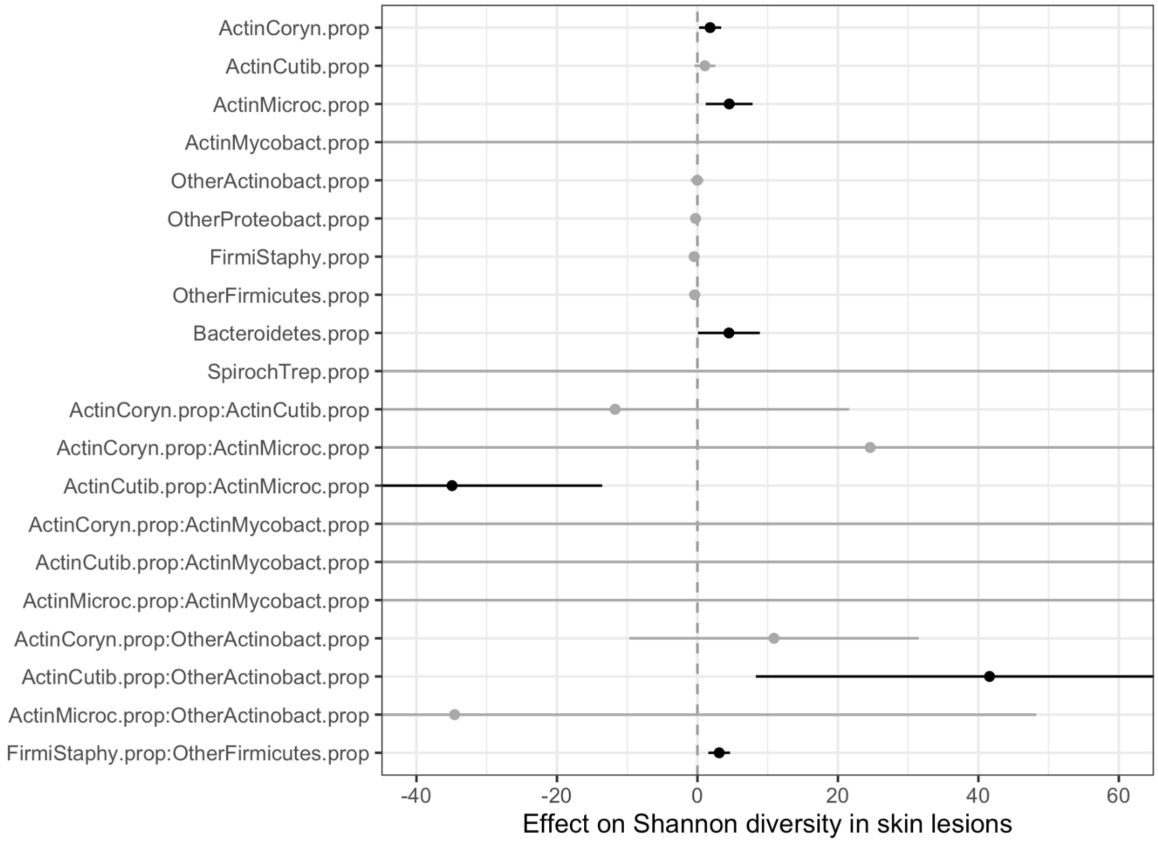


---

There is a tendency to see more reads of Corynebacteriaceae, Micrococcaceae and Bacteroidetes in samples with higher Shannon diversity. A larger proportion of *Cutibacterium* reads tends to be associated with lower Shannon diversity and fewer Micrococcaceae reads (*ActinCutib.prop:ActinMicroc.prop*). A larger proportion of *Cutibacterium* reads tends to be associated with lower Shannon diversity and fewer reads from other Actinobacteria (*ActinCutib.prop:OtherActinobact.prop*). Shannon diversity is higher when Staphylococcaceae co-occur with other Firmicutes (*FirmiStaphy.prop:OtherFirmicutes.prop*), but when one or the other is higher, lower Shannon diversity values are observed.

---

**S20. Fig . Relative abundance of taxa of interest and Shannon diversity values in skin lesions.**

The relative abundance of taxa involved in statistically significant coefficients estimated with model *sh.m284.inter.3* are shown.

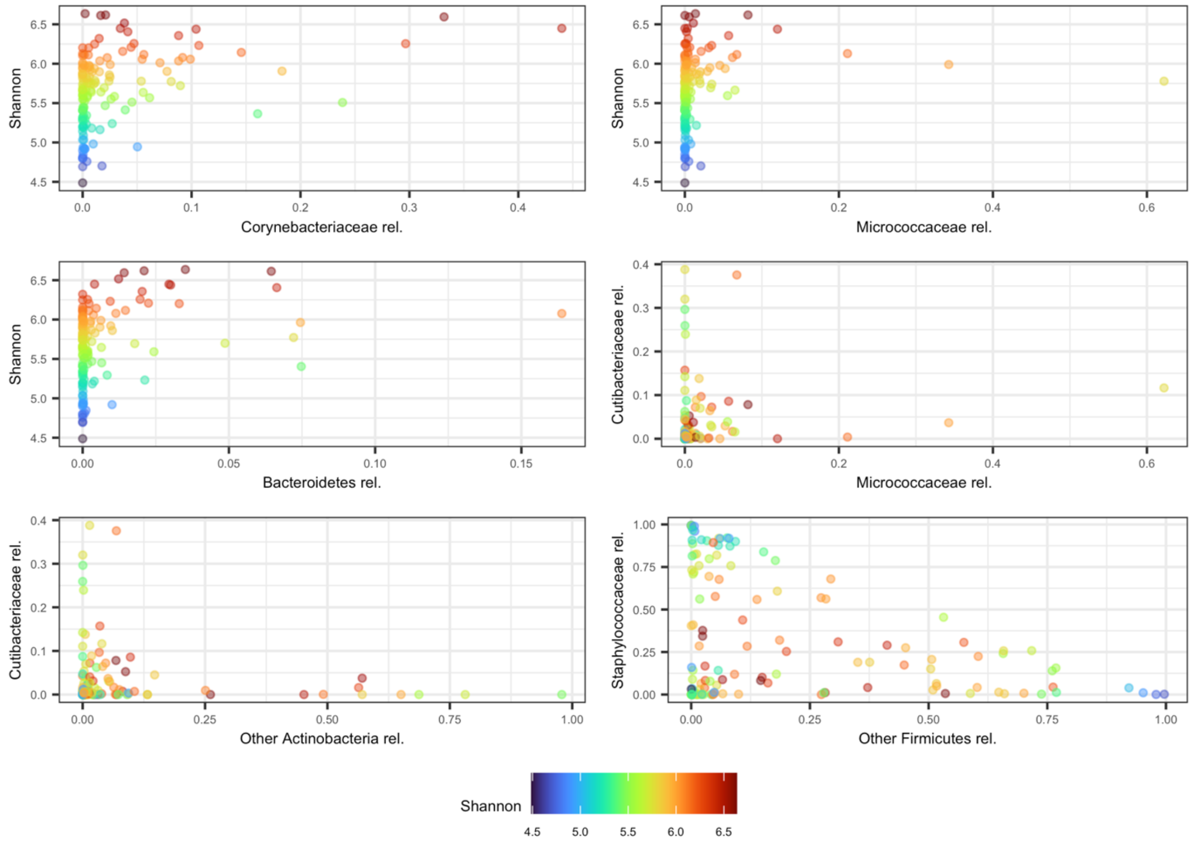


---

- Simpson diversity index

We applied the same modelling procedure to Simpson diversity values. Two simple models (*si.m1* and *si.m1v2*) were tested first and compared against a more complete model with interactions (*si.m1.inter.2*).

*si.m1: lm(Simpson ~ Lesion_location + qPCR + DPP +*
*ActinCoryn.prop +*
*ActinCutib.prop +*
*ActinMicroc.prop +*
*ActinMycobact.prop +*
*OtherActinobact.prop +*
*ProteoHaemoph.prop +*

```
          OtherProteobact.prop +
          FirmiStaphy.prop +
          OtherFirmicutes.prop +
          Bacteroidetes.prop +
          SpirochTrep.prop)
si.m1v2: lm(Simpson ~ gntl_nongntl + qPCR + DPP +
          ActinCoryn.prop +
          ActinCutib.prop +
          ActinMicroc.prop +
          ActinMycobact.prop +
          OtherActinobact.prop +
          ProteoHaemoph.prop +
          OtherProteobact.prop +
          FirmiStaphy.prop +
          OtherFirmicutes.prop +
          Bacteroidetes.prop +
          SpirochTrep.prop)
si.m1.inter.2: lm(Simpson ~ Lesion_location * gntl_nongntl +
               qPCR + DPP +
               ActinCoryn.prop *
               ActinCutib.prop *
               ActinMicroc.prop *
               ActinMycobact.prop *
               OtherActinobact.prop +
               ProteoHaemoph.prop *
               OtherProteobact.prop +
               FirmiStaphy.prop *
               OtherFirmicutes.prop +
               Bacteroidetes.prop +
               SpirochTrep.prop)
```

Here, the *anova* comparison suggested the simpler model considering the genital *vs* non-genital body sites (*si.m1v2*) captured more variance in Simpson diversity than the other two models (**S Table 4**).

**S Table 4. ANOVA comparisons of models si.m1, si.m1v2 and *si.m1.inter.2.*** These analyses of variance tables were obtained with the *anova* function comparing models *si.m1* and *si.m1.inter.2* on the one hand, and *si.m1v2* and *si.m1.inter.2* on the other hand. This shows that model *si.m1.inter.2* is significantly better than *si.m1*, but that *si.m1v2* and *si.m1.inter.2* do not significantly differ (ns). Consequently, model *si.m1v2* was favoured.

| models | Res.Df | RSS | Df | Sum of Sq | F | Pr(>F) | Signif. codes |
|---|---|---|---|---|---|---|---|
| *si.m1* | 111 | 4.91E-04 | | | | | |
| *si.m1.inter.2* | 96 | 3.73E-04 | 15 | 0.00011865 | 2.0381 | 0.01993 | * |
| *si.m1v2* | 118 | 4.98E-04 | | | | | |
| *si.m1.inter.2* | 96 | 3.73E-04 | 22 | 0.00012506 | 1.4647 | 0.1056 | ns |

We tested simpler versions of model *si.m1v2*, by removing the *gntl_nongntl*, *DPP* and *qPCR* attributes one at a time at first and then removing all three together. The pairwise *anova* comparisons to the original model (*si.m1v2*) did not suggest that these variables captured substantially more variance in Simpson diversity data than a model where they were removed. At this stage, the model retained was:

*si.m1v2nogDq: lm(Simpson ~ ActinCoryn.prop +*
*ActinCutib.prop +*
*ActinMicroc.prop +*
*ActinMycobact.prop +*
*OtherActinobact.prop +*
*ProteoHaemoph.prop +*
*OtherProteobact.prop +*
*FirmiStaphy.prop +*
*OtherFirmicutes.prop +*
*Bacteroidetes.prop +*
*SpirochTrep.prop)*

Many of the coefficients estimated by this model were not significant (the only one significant was the relative abundance of Corynebacteriaceae (*ActinCoryn.prop*). With the aim of improving the biological relevance of this model and for sake of comparison with the model retained with Shannon diversity measures, we modified this model by inserting specific interactions as follows:

*si.m2.inter: lm(Simpson ~ ActinCoryn.prop +*
*ActinCutib.prop * ActinMicroc.prop +*
*ActinMicroc.prop +*
*ActinMycobact.prop +*
*ActinCutib.prop * OtherActinobact.prop +*
*ProteoHaemoph.prop +*
*OtherProteobact.prop +*
*FirmiStaphy.prop * OtherFirmicutes.prop +*
*Bacteroidetes.prop +*
*SpirochTrep.prop)*

The formal comparison of models *si.m1v2nogDq* and *si.m2.inter* showed the latter captured significantly more variance in the data than the former (*anova* p-value < 0.001). Diagnostics statistics suggested the data point 84 may have a strong influence on the coefficients estimated by the model (**S21. Fig.**).

---

**S21. Fig. Diagnostics statistics of the model *si.m2.inter*.**

The Residuals vs Fitted plot does not show strong non-linearity, unequal error variances, or outliers as the blue line is roughly horizontal. The normal probability plot of the residuals (Normal Q-Q plot) does not suggest the residuals are not normally distributed. The Scale-Location plot does not show a clear pattern among the residuals (blue line roughly horizontal). However, the Cook's distance plot indicates specific data points may have a strong influence on the model coefficients, as distances above 0.5 are observed (especially for observation 84 – sample Y048).

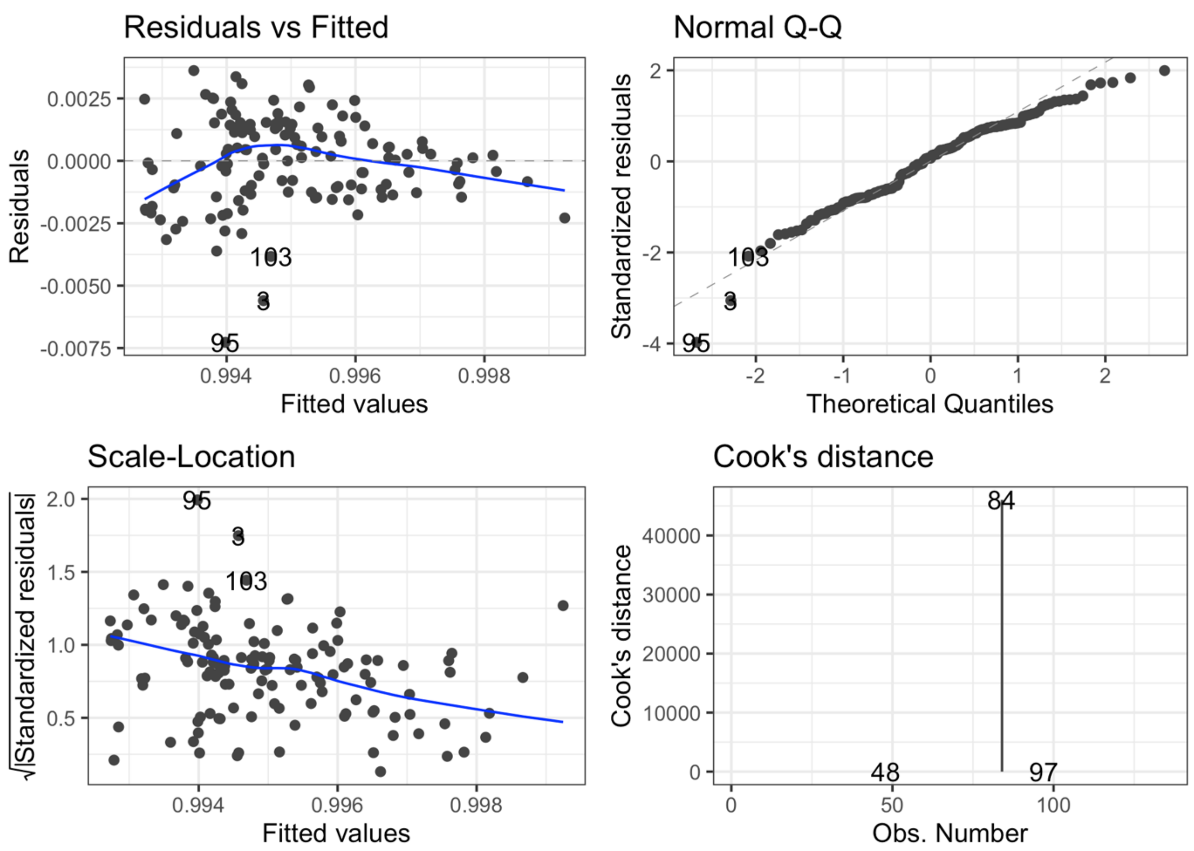


The same procedure was thus applied on a dataset containing Simpson diversity values without this data point (sample Y048). This procedure led to the same model being retained on that dataset (referred to as *si.m2.inter84* as the underlying dataset does not include sample Y048). The diagnostics statistics of this model are shown on **S22 Fig**.

**S22 Fig. Diagnostics statistics of the model *si.m2.inter84*.**

The Residuals vs Fitted plot does not show strong non-linearity, unequal error variances, or outliers as the blue line is roughly horizontal. The normal probability plot of the residuals (Normal Q-Q plot) does not suggest the residuals are not normally distributed. The Scale-Location plot does not show a clear pattern among the residuals (blue line roughly horizontal). The Cook’s distance plot does not indicate specific data points may have a strong influence on the model coefficients, as no distance above 0.5 are observed (especially since sample Y048 has been excluded from this analysis).

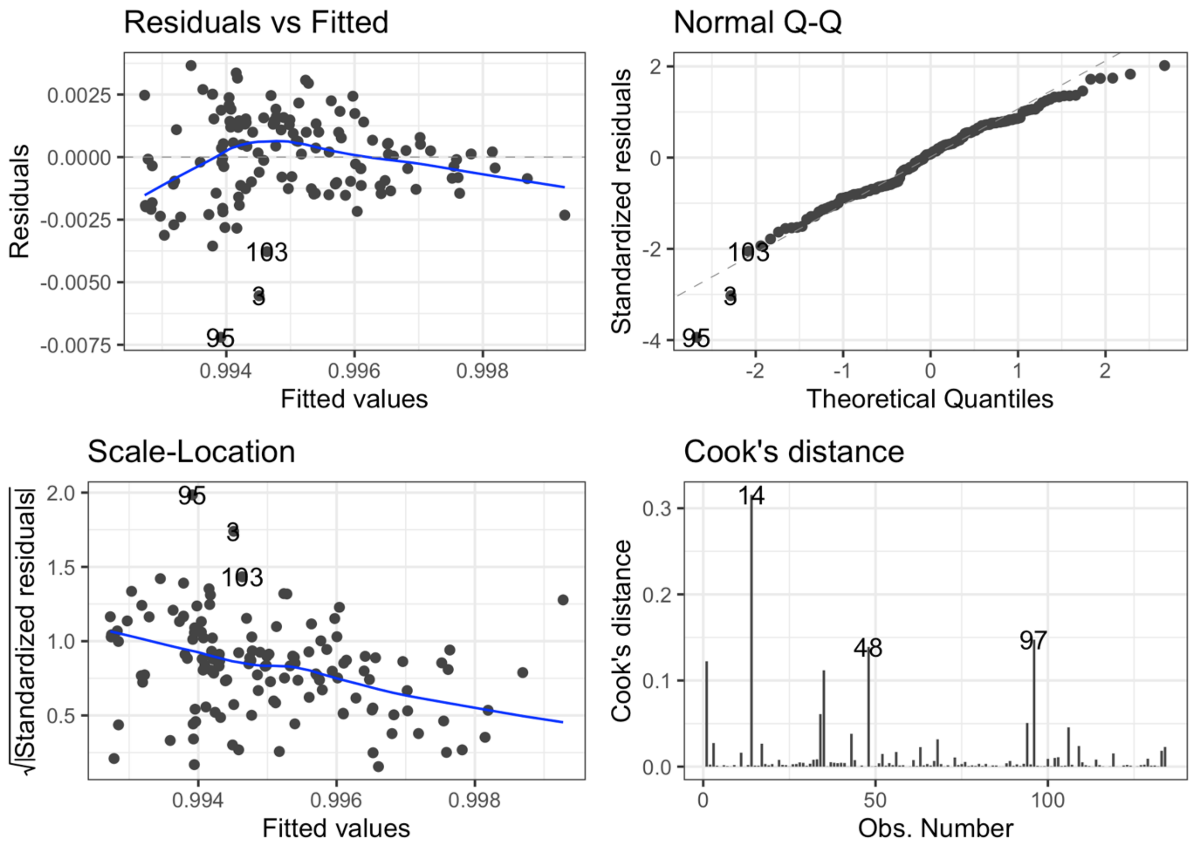


---

The coefficients estimated by this model are shown on **S Table 5**. The model retained for Shannon diversity showed six coefficients that seemed to be significantly affected by these measures, i.e. *ActinCoryn.prop*, *ActinMicroc.prop*, *Bacteroidetes.prop*, *ActinCutib.prop:ActinMicroc.prop*, *ActinCutib.prop:OtherActinobact.prop*, and *FirmiStaphy.prop:OtherFirmicutes.prop*. The model retained for Simpson diversity showed four of these seem to be significantly affected, i.e. *ActinCoryn.prop*, *ActinMicroc.prop*, *ActinCutib.prop:ActinMicroc.prop*, and *FirmiStaphy.prop:OtherFirmicutes.prop*. The direction of the effect on diversity of the relative abundance of these ubiquitous taxa is the same for these common significant coefficients.

---

**S Table 5. Coefficients estimated by the model *si.m2.inter84*.**

Coefficients in bold are those that are significantly different from 0. One coefficient could not defined because of singularities. The values provided in the column below were obtained from the *summary* of the model. Residuals: minimum (Min) and maximum (Max), and first (1Q), second (Median) and third (3Q) quantiles of the model's residuals. Estimate: estimate of the coefficient; Std. Error: residual standard error divided by the square root of the sum of the square of that particular x variable; t value: Estimate divided by Std. Error; Pr(>|t|): t value from T distribution table with the given degrees of freedom; Signif. codes: level of statistical significance, 0 '***' 0.001 '**' 0.01 '*' 0.05 '.'. For coefficients with interactions (":"), a positive estimate implies a positive effect (higher mean) of Simpson diversity on the variable to the left, and a negative estimate implies a negative effect (smaller mean) of Simpson diversity on the variable to the left.

| **Residuals** | **Min** | **1Q** | **Median** | **3Q** | **Max** |
|---|---|---|---|---|---|
| | -0.007201 | -1.15E-03 | 1.46E-04 | 0.001328 | 0.003659 |
| **Coefficients** | **Estimate** | **Std. Error** | **t value** | **Pr(>\|t\|)** | **Signif. Codes** |
| **(Intercept)** | **9.95E-01** | **1.83E-03** | **544.009** | **< 2e-16** | *** |
| **ActinCoryn.prop** | **8.48E-03** | **3.08E-03** | **2.751** | **0.00686** | ** |
| ActinCutib.prop | 5.34E-03 | 3.33E-03 | 1.604 | 0.1114 | |
| **ActinMicroc.prop** | **1.79E-02** | **5.46E-03** | **3.283** | **0.00134** | ** |
| ActinMycobact.prop | 1.15E-01 | 1.80E-01 | 0.636 | 0.52579 | |
| OtherActinobact.prop | 1.06E-03 | 2.21E-03 | 0.478 | 0.63381 | |
| ProteoHaemoph.prop | NA | NA | NA | NA | |
| OtherProteobact.prop | -1.06E-03 | 1.86E-03 | -0.573 | 0.56788 | |
| FirmiStaphy.prop | -2.29E-03 | 1.88E-03 | -1.218 | 0.22575 | |
| OtherFirmicutes.prop | -2.02E-03 | 1.96E-03 | -1.03 | 0.30497 | |
| Bacteroidetes.prop | 1.37E-02 | 1.11E-02 | 1.239 | 0.21766 | |
| SpirochTrep.prop | 6.12E+00 | 6.00E+00 | 1.019 | 0.31037 | |
| **ActinCutib.prop:ActinMicroc.prop** | **-1.62E-01** | **5.36E-02** | **-3.029** | **0.003** | ** |
| ActinCutib.prop:OtherActinobact.prop | 1.25E-01 | 6.31E-02 | 1.977 | 0.05031 | |
| **FirmiStaphy.prop:OtherFirmicutes.prop** | **1.69E-02** | **3.86E-03** | **4.37** | **2.66E-05** | *** |

Residual standard error: 0.001869 on 120 degrees of freedom
Multiple R-squared: 0.3772, Adjusted R-squared: 0.3097
F-statistic: 5.59 on 13 and 120 DF, p-value: 7.018e-08

---

The coefficients estimated by this model are presented in **S23 Fig**. These coefficients can be better appreciated when placed in the context of the underlying diversity (**S24 Fig.**).

---

**S23 Fig. Dot-and-Whisker plot of *si.m2.inter84* regression model results.** The whiskers show the 95% confidence interval of the coefficient estimates. The x-axis is scaled to provide an optimal overview of the statistically significant (black) and not statistically significant (grey) coefficients. The dashed vertical line shows no effect on Simpson diversity.

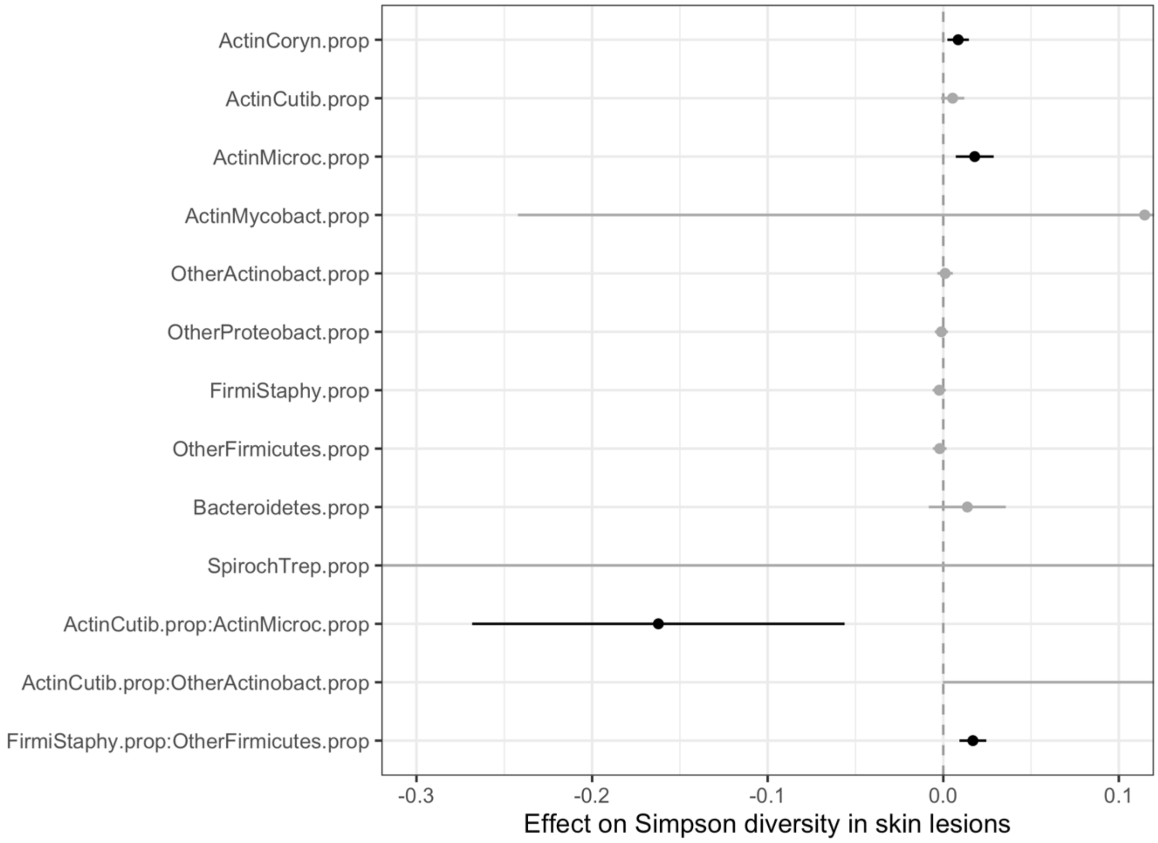


---

There is a tendency to see more reads of Corynebacteriaceae and of Micrococcaceae in samples with higher Simpson diversity. A higher Simpson diversity tends to be observed when both the relative abundance of *Cutibacterium* and those of Micrococcaceae are intermediate (*ActinCutib.prop:ActinMicroc.prop*). However, when the relative abundance of *Cutibacterium* is high, there tend to be fewer Micrococcaceae. A higher Simpson diversity tends to be observed when both the number of reads of Staphylococcaceae and those of other Firmicutes are intermediate, when one or the other is very high, Simpson diversity tends to be low (*FirmiStaphy.prop:OtherFirmicutes.prop).* As the terms that were significantly associated with the values of Simpson diversity were also significantly associated with the values of Shannon diversity, and were going in the same directions, these are presented together in the main text (**Fig. 7**).

**S24. Fig. Relative abundance of taxa of interest and Simpson diversity values in skin lesions.**

The relative abundance of taxa involved in statistically significant coefficients estimated with model *si.m2.inter84* are shown.

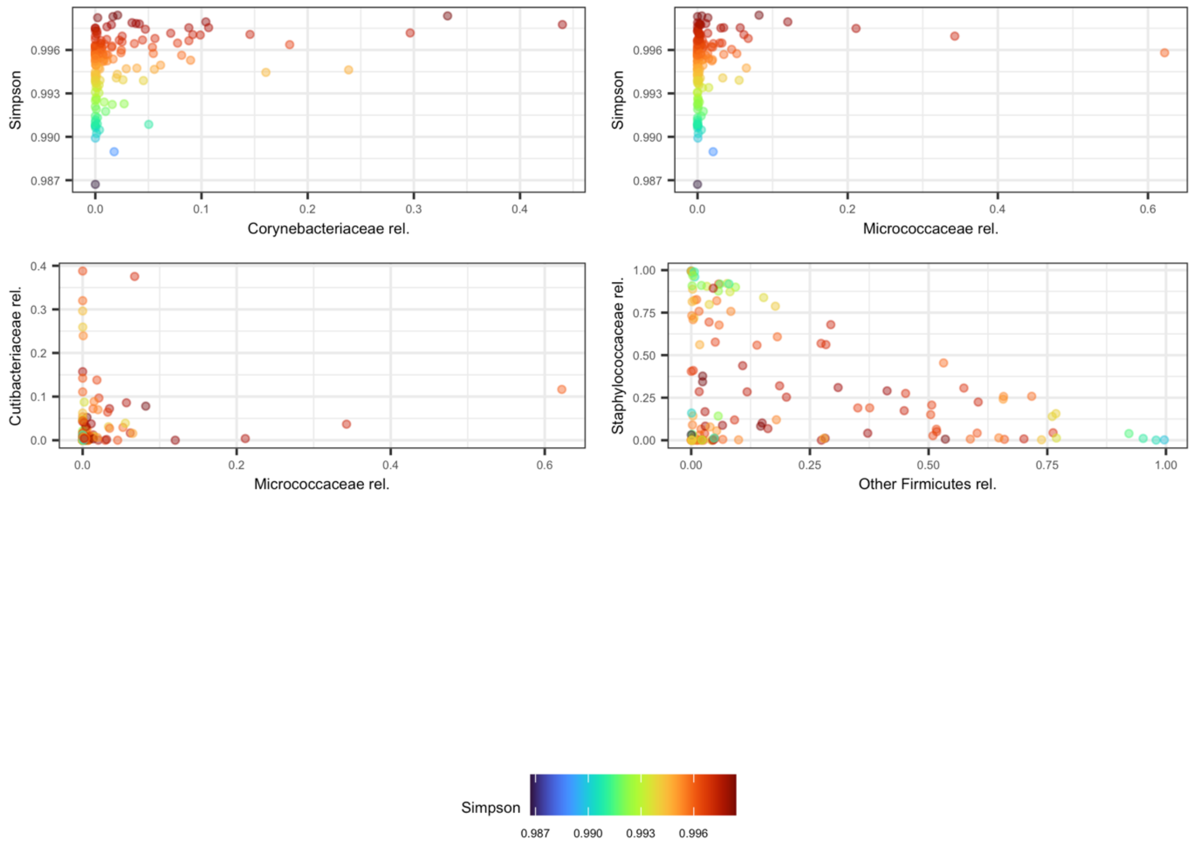


We warrant against using these linear models for predictive purposes. Linear modelling is used here as a means to investigate association in the data in a statistically robust manner as they embed some corrections for multiple testing. We believe they are useful approaches to formulate hypotheses that need to be investigated further with distinct datasets.
